# Mutation of *IDR1* enhances drought tolerance by reducing ROS production and activating ROS scavenging in rice

**DOI:** 10.1101/2020.08.24.264556

**Authors:** Xiaofeng Zu, Yanke Lu, Qianqian Wang, Yumei La, Feng Tan, Jiayu Niu, Huihui Xia, Xinyue Hong, Yufeng Wu, Shaoxia Zhou, Kun Li, Huhui Chen, Sheng Qiang, Qi Rui, Huaqi Wang, Honggui La

**Author notes:** Corresponding author: Honggui La, College of Life Sciences, Nanjing Agricultural University, Nanjing 210095, China. Contributed equally to the article (Xiaofeng Zu, Yanke Lu, Qianqian Wang). **Author contributions** H. L, X. Z and Y. L conceived and designed the original research plans; H. L. supervised the entire experiments; X. Z., Y. L. and Q. W. performed most of the experiments; Y. L. analyzed the mRNA-seq data; F. T., J. N., H. X., X. H., K. L. and H. C. helped prepare samples; Y. W., S. Z., S. Q. and Q. R. provided analytical and experimental assistance; H. L., X. Z., Y. L. and Q. W. wrote the article; H. L. and H. W. revised the manuscript. **Responsibilities of the author for contact** Honggui La. **Email address of author for contact** Xiaofeng Zu, Yanke Lu, Qianqian Wang, Yumei La, Feng Tan, Jiayu Niu, Huihui Xia, Xinyue Hong, Yufeng Wu, Shaoxia Zhou, Kun, Li, Huhui Chen, Sheng Qiang, Qi Rui, Huaqi Wang.

## Abstract

To discover new mutant alleles conferring enhanced tolerance to drought stress, we screened a mutagenized rice population (cv. IAPAR9) and identified a mutant, named *idr1-1* (*for increased drought resistance 1-1*), with obviously increased drought tolerance under upland field conditions. The *idr1-1* mutant possessed a significantly enhanced ability to tolerate high-drought stress in different trials. Map-based cloning revealed that the gene LOC_Os05g26890 (corresponding to *D1* or *RGA1* gene), residing in the mapping region of *IDR1* locus, carried a single-base deletion in the *idr1-1* mutant, which caused a frameshift and premature translation termination. Complementation tests indicated that such a mutation was indeed responsible for the elevated drought tolerance in *idr1-1* mutant. IDR1 protein was localized in nucleus and to plasma membrane or cell periphery. Further investigations indicated that the significantly increased drought tolerance in *idr1-1* mutant stemmed from a range of physiological and morphological changes occurring in such a mutant, including greater leaf potentials, increased proline contents, heightened leaf thickness, and upregulation of antioxidant-synthesizing and drought-induced genes, etc., under drought-stressed conditions. Especially, ROS production from NADPH oxidases and chloroplasts might be remarkably impaired, while ROS-scavenging ability appeared to be markedly enhanced as a result of significantly elevated expression of a dozen ROS-scavenging enzyme genes in *idr1-1* mutant under drought-stressed conditions. Besides, IDR1 physically interacted with TUD1, and *idr1-1* mutant showed impaired EBR responsiveness. Altogether, these results suggest that mutation of IDR1 leads to alterations of multiple layers of regulations, which ultimately confers obviously enhanced drought tolerance to the *idr1-1* mutant.

**One-sentence summary:** Mutation of *IDR1* significantly enhances drought tolerance in an upland cultivar IAPAR9 by decreasing apoplastic and chloroplastic ROS production and increasing ROS-scavenging ability

## Introduction

Rice is one of the most important crops worldwide. Mainly due to the increasing shortage of water resources, drought has become the major limiting factor for rice growth and production. Hence, breeding and cultivation of drought-tolerance rice varieties are important measures to cope with outcomes of drought stress. Thus, understanding the mechanisms of how rice plants respond to drought stress and what factors play critical role in contributing to drought tolerance has become an important step towards breeding drought-tolerant rice by means of conventional cross and genetic engineering.

Over the past decades, much progress had been made in understanding the mechanisms of how drought stress damages plants and how plants defend against the drought stress to safeguard themselves from severe damage. Plant stomata are mainly responsible for water transpiration and gas exchange. Stomatal movements regulate water status in response to drought stress (Hong et al., 2008). Phytohormone abscisic acid (ABA) performs important functions in modulating stomata movements when plants are subjected to drought stress (Tuteja and Sopory, 2008). It is generally known that stomatal closing regulated by ABA plays essential roles in preventing plants from excessive water loss at the early stages of drought stress. Besides, other phytohormones, like GA, brassinosteroid (BR), ethylene, auxin, etc., are all involved in plant drought responses (Desikan et al., 2006; Ge et al., 2015; Shi et al., 2015). Growing evidence demonstrates that disruption of GA signaling pathway brings about transcriptional elevation of *GA2ox*, a gene encoding GA-inactivating enzyme, and *RGL3* that encodes a DELLA protein, eventually leading to plant growth retardation and enhanced drought tolerance (Colebrook et al., 2014). DELLA protein was found to restrain the accumulation of ROS which is usually overaccumulated when plants are exposed to abiotic stresses. BRs are a class of polyhydroxylated sterol derivatives, and they play vital roles in promoting cell expansion and division, regulating senescence, and modulating plant responses to various environmental cues (Hu et al., 2013; Feng et al., 2015). A recent report showed that 24-epibrassinolide (EBR) induced stomatal closure in Arabidopsis by triggering a signal transduction pathway including ethylene synthesis, activation of Gα protein, and hydrogen peroxide (H_2_O_2_) and nitric oxide (NO) production (Shi et al., 2015). Another study reported that BR was able to induce H_2_O_2_ production, and the H_2_O_2_ played vital roles in BR signaling (Tian et al., 2018). In *Brachypodium distachyon*, *BRASSINOSTEROID INSENSITIVE 1* (*BdBRI1*) RNAi lines exhibited markedly enhanced drought tolerance, accompanied by highly elevated expression of drought-responsive genes, such as *BdCOR47*/*BdRD17*, *BdERD1* and *BdRD26* (Feng et al., 2015). Arabidopsis *wrky46 wrky54 wrky70* triple mutant had defects in BR-regulated growth and was more tolerant of drought stress, suggesting that WRKY46/54/70 is positively involved in BR-regulated growth and negatively involved in the drought responses (Chen et al., 2017).

Accumulating evidence shows that stomatal closure is just an early response when drought stress occurs (Distefano et al., 2015). If the stress becomes severe, plants activate other mechanisms to protect proteins and membrane structures in order to avoid cell death caused by water deficit, such as accumulation of cuticular wax crystals and prolines, control of ROS homeostasis, as well as transcriptional activation of large numbers of transcription factors (Distefano et al., 2015). The rice *drought-induced wax accumulation 1* (*dwa1*) mutant was impaired in cuticular wax accumulation under drought stress, which resulted in significantly increased drought sensitivity (Zhu and Xiong, 2013). Proline is thought to be a scavenger of free radicals and plays an important role in enhancing drought tolerance via osmotic adjustment. In *BdBRI1* RNAi lines, expression of *BdP5CS* was highly induced in response to drought stress, which might be one of the reasons for significantly enhanced drought tolerance in such RNAi lines (Feng et al., 2015).

Production of ROS is a frequent event in plants that suffer from diverse abiotic stresses. There are many potential sources of ROS in plant cells, including chloroplasts, mitochondria, peroxisomes, plasma membrane NADPH oxidases, cell wall peroxidases, apoplastic oxalate oxidases, and amine oxidases (Zhang et al., 2010). Mounting evidence showed that ROS generated by NADPH oxidases play crucial roles in plant abiotic stresses, defense and hormonal responses. However, excessive amounts of ROS are harmful to the plants because it causes damage to cells through oxidizing proteins, lipids, and DNA, ultimately leading to retarded growth and eventual cell death (Bargmann and Munnik, 2006; Ning et al., 2010; You et al., 2014). ROS is also able to cause photoinhibition occurring in PSII reaction center, which, in turn, triggers undue production of the ROS (Ferrero-Serrano et al., 2018). Thus, to counteract the ROS’s deleterious effects, plants have evolved the mechanisms for scavenging these excessive ROS, including the involvement of a variety of ROS-scavenging enzymes, superoxide dismutase (SOD), catalase (CAT), glutathione reductase, and peroxidase (POX), etc. (You et al., 2014). Mutation of ROS scavenger’s genes reportedly leads to drought sensitivity in plants. For example, a *dsm1* mutant defective in a putative MAPK kinase kinase gene in rice, showed more sensitive to oxidative stresses due to an increase in ROS level caused by reduced expression of *POXs* (*OsPOX22.3* and *OsPOX8.1*) (Ning et al., 2010). The *ospp18* mutants exhibited increased sensitivity to oxidative stress because several genes encoding ROS-scavenging enzymes were downregulated in such mutant plants (You et al., 2014). Overexpression (OE) of *OsGH3-2* in rice plants decreased free IAA level and thus impaired SOD activity, therefore conferring drought sensitivity to such plants (Du et al., 2012). OsLG3, a ERF family transcription factor, played a positive role in drought stress tolerance in rice by inducing ROS scavenging; overexpression of the *OsLG3* could reduce overaccumulation of ROS as a total of 10 ROS-scavenging genes (*APXs*, *CATB*, *PODs*, *SODs*, etc.) were upregulated in the OE lines (Bargmann and Munnik, 2006).

Heterotrimeric GTP-binding proteins (G proteins), comprising three subunits Gα, Gβ and Gγ, are key signaling molecules in all eukaryotes. Classically, signal perception by a G protein-coupled receptor results in the dissociation of Gα from Gβγ subunits, and then Gα or Gβγ dimer can separately activate downstream signaling (Ferrero-Serrano and Assmann, 2016). Arabidopsis genome has only one Gα (*GPA1*), one Gβ (*AGB1*), and three Gγ (*AGG1* to *AGG3*) genes; rice genome contains one Gα (*RGA1*), one Gβ (*RGB1*), four Gγ subunit (*RGG1*, *RGG2*, *GS3*, and *DEP1*), and one Gγ-related gene (or pseudogene) *OsGGC2* (Ferrero-Serrano and Assmann, 2016). G proteins play vital roles in the transduction of extracellular signals into intracellular responses; in plants, they have been implicated in multiple signaling pathways, including phytohormone responses such as GA, ABA, jasmonic acid, auxin, and BR (Ritchie and Gilroy, 2000; Wang et al., 2001; Wang et al., 2006; Oki et al., 2009; Hu et al., 2013; Ge et al., 2015; Shi et al., 2015; Ferrero-Serrano and Assmann, 2016). In Arabidopsis, the GPA1 is a modulator of plant cell proliferation, because *gpa1* null mutant has reduced cell division in aerial tissues throughout development (Ullah et al., 2001). The GPA1 also regulates stomatal density by controlling epidermal cell sizes and stomata formation (Nilson and Assmann, 2010). Importantly, G proteins not only participate in stomatal movements, but also act as key regulators of H_2_O_2_ production in guard cells induced by ABA (Ge et al., 2015). Furthermore, *gpa1* mutation resulted in insensitivity of stomatal opening to inhibition triggered by ABA, but did not cause significant impairment of ABA-induced stomatal closing, with the result that the rates of water loss from *gpa1* mutants were greater than those from the wild-type control plants (Wang et al., 2001). In *gpa1-3* and *gpa1-4* guard cells, ABA failed to induce ROS production, suggesting that the GPA1 acts upstream of ROS production in the ABA signaling pathway (Zhang et al., 2011). The *gpa1* mutants also showed defects in ethylene-induced H_2_O_2_ production and stomatal closure, whereas wGα and cGα OE lines showed faster stomatal closure and greater H_2_O_2_ production in response to ethylene, indicating that Gα mediates ethylene-induced stomatal closure via H_2_O_2_ production (Ge et al., 2015). Moreover, *gpa1-4* leaves displayed less damage when treated with O_3_, and O_3_-induced H_2_O_2_ was found to be greatly impaired in the mutant leaves, demonstrating that the resistance of the *gpa1-4* mutants to O_3_-induced cell damage is attributable to the defect in Gα-mediated activation of cell death-associated late component of the oxidative burst (Joo et al., 2005). Likewise, cell death of *gpa1* was mitigated as well in phyA-mediated cell-death pathway (Wei et al., 2008). In addition, *gpa1* showed approximately 100-fold less responsiveness to GA (Ullah et al., 2002), and reduced responsiveness to brassinolide (BL), suggesting that GA and BR signal transduction is likely coupled by the Gα protein (Ullah et al., 2001; Ullah et al., 2002).

The rice Gα gene (named *D1* or *RGA1*) was first identified from a rice *Dwarf 1* (*D1*) mutant by using a map-based cloning strategy (Ashikari et al., 1999). *d1* mutant showed a dwarf phenotype; the elongation of the second leaf sheaths of *d1* seedlings was much less responsive to exogenous GA_3_ than that of the wild-type seedlings, indicating that the *d1* mutation partially impairs GA-signaling pathway (Ueguchi-Tanaka et al., 2000). Two independent studies all indicated that D1 participated in BR responses, and the *d1* mutant displayed a remarkably reduced responsiveness to EBR, suggesting that the D1 is involved in BR signaling (Wang et al., 2006; Oki et al., 2009). Further, a D1-interacting protein Taihu Dwarf1 (TUD1), a U-box E3 ubiquitin ligase, was reported to act in conjunction with the D1 to affect the BR-signaling pathway, because *tud1* mutant was also less responsive to BR treatment as was the *d1* mutant (Hu et al., 2013; Ferrero-Serrano and Assmann, 2016). Another study provided a line of evidence that epidermal cell death in rice was dependent on signaling through Gα proteins (Steffens and Sauter, 2009). A recent study demonstrated that *d1* mutants exhibited significantly reduced sensitivity to drought stress; after a long-term progressive drought stress, the *d1* plants showed obviously more tolerance to the stress than wild-type plants (Ferrero-Serrano and Assmann, 2016). Moreover, during the course of drought stress, the decrease in photosynthetic rates in the *d1* mutants was markedly slower compared to the control plants, implicating that limitation of photosynthesis caused by drought are less severe in the *d1* mutants (Ferrero-Serrano and Assmann, 2016). Additionally, under NaCl-treated conditions the *d1* mutants displayed remarkable alleviation of leaf senescence, chlorophyll degradation, and cytoplasm electrolyte leakage (Urano et al., 2014).

In this study, we identified a highly drought-tolerant rice mutant in upland fields, named *idr1-1*, by screening a rice population undergoing ^60^Co γ-ray radiation. *idr1-1* mutants consistently showed obviously enhanced drought tolerance under different growth conditions; map-based cloning revealed that the *idr1-1* was a new mutant allele of Gα. Detailed investigations demonstrated that the enhanced drought tolerance of *idr1-1* mutant appeared to stem from altered morphology, impaired ROS production, increased ROS scavenging, weakened leaf senescence, elevated antioxidant levels, etc.; in addition, IDR1 physically interacted with TUD1, and the *idr1-1* mutant showed reduced BR responsiveness, which is probably associated with the enhanced drought tolerance of *idr1-1* mutant. Given the significantly enhanced drought tolerance caused by the *idr1-1* mutation, the *idr1-1* mutant allele has considerable potential for rice drought-tolerant breeding.

## Results

### Identification of an *idr1-1* Mutant that Shows Significantly Enhanced Drought Tolerance in Upland Field Conditions

To study the mechanisms of rice drought tolerance, we performed a genetic screening in upland fields to find candidate lines with enhanced drought tolerance from a γ-ray-mutagenized upland rice (cv. IAPAR9) population. Eventually, a candidate mutant plant was discovered to have obviously increased drought tolerance, and was thus referred to as *idr1-1* (for *increased drought resistance 1-1*). The drought-tolerant phenotype of *idr1-1* mutant was further reconfirmed in pots and concrete tanks, respectively (Fig. 1, A and C). For drought treatments in pots, the *idr1-1* mutants exhibited enhanced drought tolerance to severe drought stress compared to IAPAR9 (Fig. 1A), and the survival rate of *idr1-1* mutants following rewatering was approximately 64% higher than that of IAPAR-9 plants (Fig. 1B). Further, for drought treatments in concrete tanks, under moderate drought stress conditions, IAPAR9 plants became severely wilted, whereas *idr1-1* mutants grew quite normally (Fig. 1C). Under severe drought stress conditions, most of IAPAR9 leaves became dry while only a quite small fraction of *idr1-1* leaves became wilted or dry. After rewatering, about 50% of IAPAR9 plants undergoing moderate drought stress died but more than 95% of *idr1-1* mutants still stay alive (Fig. 1D). Except for drought-tolerant phenotype, *idr1-1* mutants also displayed other developmental phenotypes, such as noticeable dwarfism, shorter lengths of grains as well as panicles and leaves, compared to IAPAR9 plants (Supplemental Fig. S1). However, grain width of the *idr1-1* mutants exhibited no difference from that of the IAPAR9 plants (Supplemental Fig. S1E, right panel). When germinated in the presence of different concentrations of glucoses, *idr1-1* seeds showed significantly reduced germination rates under the conditions of 6% and 8% glucoses (Supplemental Fig. S2, A and B), suggesting that the *idr1-1* mutants have increased sensitivity to high concentrations of glucoses, which is in agreement with the previous observations in Arabidopsis (Ritchie and Gilroy, 2000). When grown in 150 mM NaCl solution, *idr1-1* mutants appeared higher sensitivity than IAPAR9 plants as approximately 10% of mutant plants and 47% of IAPAR9 plants remained alive following the treatments (Supplemental Fig. S2, C and D). Taken together, these results demonstrate that mutation of *IDR1* leads to a wide range of morphological and physiological changes, and the *idr1-1* mutants show obviously increased drought tolerance under different growth conditions.

**Figure 1.**
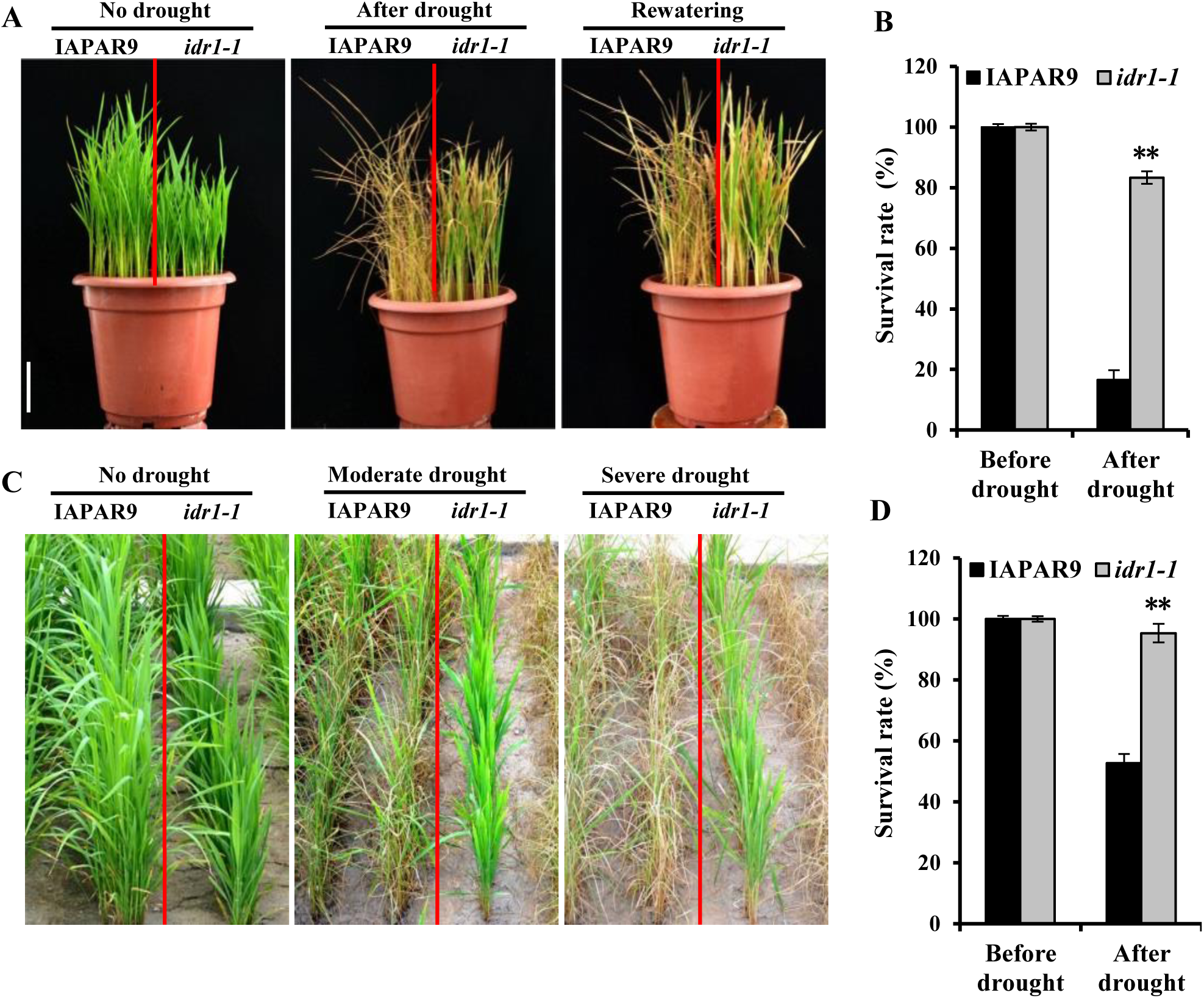
*idr1-1* mutation confers obviously enhanced drought tolerance to upland rice plants of cv. IAPAR9 under different growth conditions. **(A and B)** Drought tolerance tests of *idr1-1* mutant and IAPAR9 seedlings in pots. Both the *idr1-1* mutant and IAPAR9 seedlings at a V6 (when the sixth leaf has a visible “collar” at the base of the leaf) developmental stage were subjected to drought stress for approximately 2-3 weeks (severe drought), followed by rewatering for 7 days to examine their drought tolerance and calculate survival rates (A). Survival rates of *idr1-1* mutant and IAPAR9 seedlings as described in A were calculated from rewatered plants (B). Scale bar = 5 cm. **(C and D)** Drought tolerance tests of *idr1-1* mutant and IAPAR9 seedlings in concrete tanks. Both *idr1-1* mutant and IAPAR9 seedlings at the V6 developmental stage grown in concrete tanks with a rain-off shelter were subjected to moderate (withholding water for approximately 2 weeks) or severe drought stress (withhold water for approximately 3 weeks) to examine their drought tolerance (C). Survival rates of *idr1-1* mutant and IAPAR9 plants were calculated from rewatered plants from the experiment depicted in C (D). The pictures were taken after drought treatment. Data are means (±SD) for three biological replicates. Significant differences from IAPAR9 were determined by Student’s *t*-test: **P < 0.01, t-test.

### Map-Based Cloning of *IDR1* Gene

F1 heterozygous plants produced from a cross between *idr1-1* mutant female and IAPAR9 male showed similar performance of drought tolerance as the IAPAR9 plants, and were noticeably less tolerant of drought stress compared with the *idr1-1* mutants (Supplemental Fig. S3A), suggesting that *idr1-1* mutation is recessive. For cloning *IDR1* gene, we crossed the *idr1-1* mutant females separately with each of 10 upland rice cultivar males, including HD297, HD385, HD119, etc. (Supplemental Table S1). The resulting F2 populations were planted in the above-mentioned concrete tanks for the moderate drought treatment to distinguish drought-tolerant and drought-sensitive individuals. The results indicated that the F2 progeny produced by a cross between *idr1-1* mutant and HD119 plant yielded a segregation of 160 highly tolerant and 522 mildly tolerant plus sensitive plants (M+S) (in a ratio of 1:3.2), which is in good agreement with the expected 1:3 ratio (X^2^ = 2.01, P < 0.05) (Supplemental Fig. S3; Supplemental Table S1), suggesting that the *idr1-1* mutation are recessive. Furthermore, we chose 160 F2 individuals with highly drought tolerance from such a population for the primary mapping. Eventually we mapped the candidate mutation to a ∼0.78Mb genomic interval between markers RM18402 and RM18421 on chromosome 5 (Fig. 2A). Further, 3800 F2 individuals were used for narrowing down the candidate region. Finally, *IDR1* was delimited to an interval of ∼30.7 kb between markers IM63 and IM59, which included four predicted genes (Fig. 2A). After sequencing, a single-base deletion (position 886 downstream of translation initiation codon ATG) in the fifth exon of LOC_Os05g26890 (corresponding to *D1* or *RGA1* gene) in *idr1-1* mutant was identified, which as a result led to a truncated protein consequent upon a premature stop codon (TGA) newly created on the open reading frame (ORF) of *D1* (Fig. 2B) (Ashikari et al., 1999; Wang et al., 2006). The *D1* had been previously reported to encode α subunit of heterotrimeric G proteins (also known as *RGA1*) (Wang et al., 2006; Oki et al., 2009); nevertheless, there were very few reports on the involvement of D1 in rice drought tolerance.

**Figure 2.**
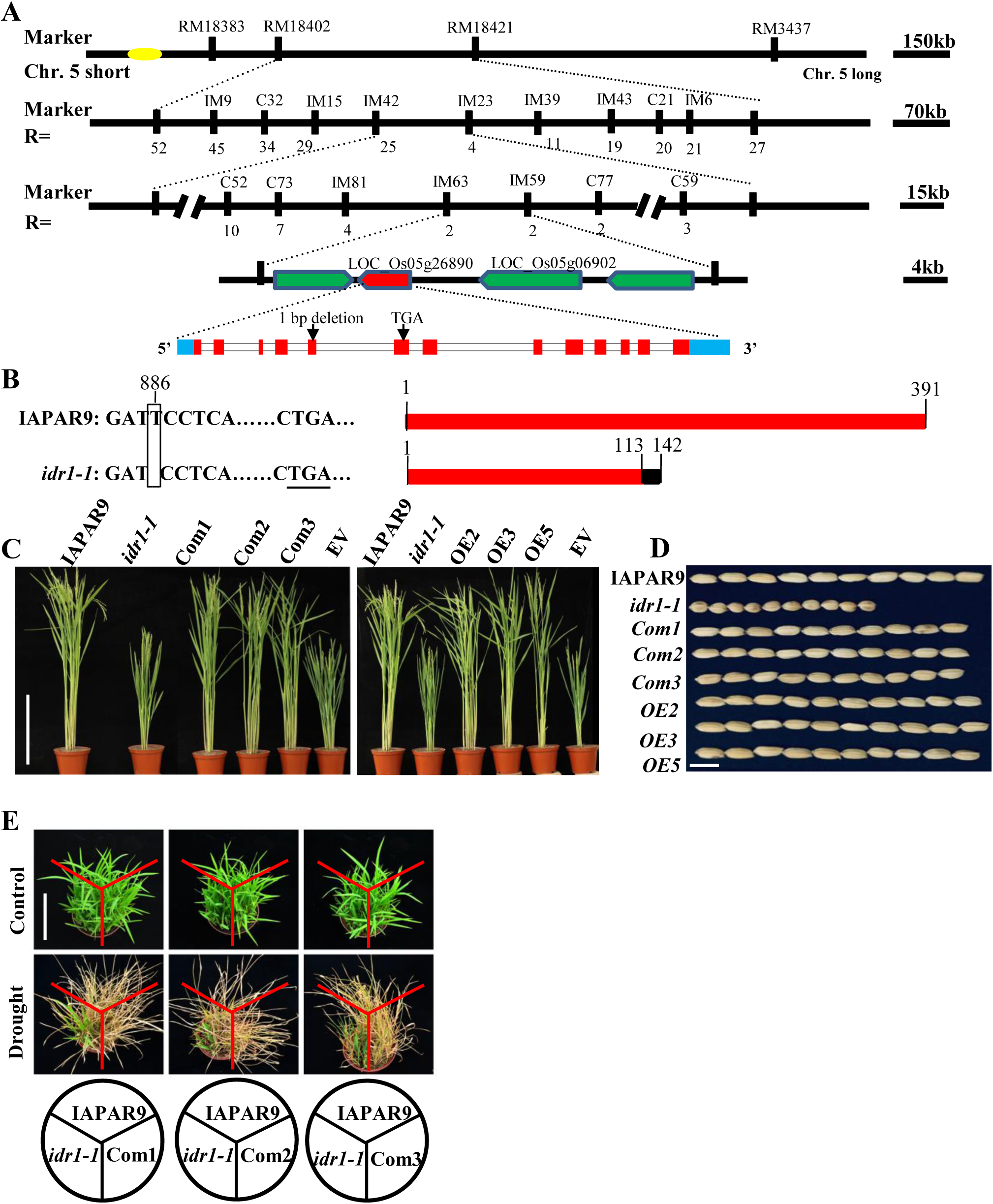
Map-based cloning of *IDR1* and complementation of *idr1-1* mutant by *IDR1* transgenes. **(A)** Mapping of *IDR1* locus on the chromosome 5 by using molecular markers. *IDR1* locus was primarily mapped to the interval between molecular markers RM18402 and RM18421 on the chromosome 5, and it was further delimited to a ∼30.7 kb genomic region between markers IM63 and IM59. Marker names and the numbers of recombinants were shown above and below each line, respectively. Four candidate genes were indicated by arrows, among which the red arrow represented the *IDR1* gene (LOC_05g26890). The two genes without locus names above them were predicted by RAP-DB database but not by MSU RGAP database. A single-base deletion occurring on the fifth exon of *IDR1* was shown, whereby a premature stop codon (TGA) was created on the ORF sequence of such a gene. **(B)** A close-up view of the single-base deletion at the position 886 downstream of the ATG start codon on the *IDR1* genomic DNA (left panel). The newly created premature stop codon TGA on the *IDR1* ORF sequence was underlined. The full-length and truncated proteins corresponding to the entire ORF sequence and the mutated ORF sequence of *IDR1* were shown in right panel. Black bar represents a portion of protein with a changed sequence due to the frameshift resulting from the single-base deletion. The numbers above the diagrams indicate the coordinates relative to the start codon ATG (left panel) or start amino acid 1 (right panel). **(C and D)** Reversion of plant heights (C) and grain lengths (D) of *idr1-1* mutants to wild-type sizes by introduction of *IDR1* genomic DNA or OE constructs to the mutants. Com1 to Com3, *IDR1*

Complementation of the mutant with a wild-type *IDR1* genomic DNA fragment completely corrected the dwarfism defect of *idr1-1* mutants (Fig. 2C); by contrast, the transformed plants carrying empty vector (EV) bore striking similarity in plant height to the *idr1-1* mutants (Fig. 2C). Similarly, overexpression of *IDR1* cDNA in the *idr1-1* mutant background also totally rescued the *idr1-1*’s dwarfism phenotype (Fig. 2C). Moreover, the seeds harvested from *IDR1* complementation lines or OE lines reverted to approximately their wild-type sizes (Fig. 2D). As expected, the complementation lines grown in pots exhibited obviously reduced drought tolerance, which was comparable to the performance as was observed in IAPAR9 plants (Fig. 2E); that is, both wild-type IAPAR9 plants and *IDR1* complementation lines were almost completely dead, while a fraction of *idr1-1* mutants remained alive (Fig. 2E). Likewise, when grown in the concrete tanks, such complementation lines displayed remarkable sensitivity to moderate and severe drought treatments (Supplemental Fig. S4). Hence, it can be concluded from the aforementioned results that reintroduction of *IDR1* into the *idr1* mutants does indeed lower the drought tolerance of the *idr1-1* mutants to close to the wild-type level. When all of the genotypes (which underwent severe drought stress) grown in the above-mentioned concrete tanks grew to maturity, the average seed setting rate of *idr1-1* mutant plants was approximately 40%, which was around 2 times as high as that of IAPAR9 plants (which achieved about 20% average seed setting rate) (Fig. S5). This suggests that loss of function of IDR1 is able to reduce loss of grain yield under the conditions of severe drought stress.

### Expression Patterns and Subcellular Localization of IDR1

Expression analyses revealed that transcript levels of *IDR1* gradually declined alongside the increasing drought stresses (from control to severe drought stress), which points to the possibility that the expression of *IDR1* is suppressed by drought stress (Fig. 3A). Twenty percent PEG treatments also led to significant drop in *IDR1* expression to around half of the level prior to treatment (0 h), at 4 and 8 h following treatments (Fig. 3B). By contrast, 150 mM NaCl treatments caused continuous induction of *IDR1* expression during the first 4 hours after treatments, followed by a reduction to the level similar to that at 0 h (Fig. 3C). Cold treatments (4°C) brought about weak upregulation of *IDR1* expression at 1 h upon treatment, but the expression was downregulated from 4 hour onwards (Fig. 3D); similarly, *IDR1* expression was also induced by heat treatments (40°C), peaking at 4 h, declining at 8 h (Fig. 3E). Fifty micromolar ABA treatments made *IDR1* downregulated, which is similar to the case in the foregoing drought treatments (Fig. 3, F and A). Altogether, these data suggest that *IDR1* expression is affected by different types of stresses and downregulation of *IDR1* may contribute to drought tolerance of rice plants.

**Figure 3.**
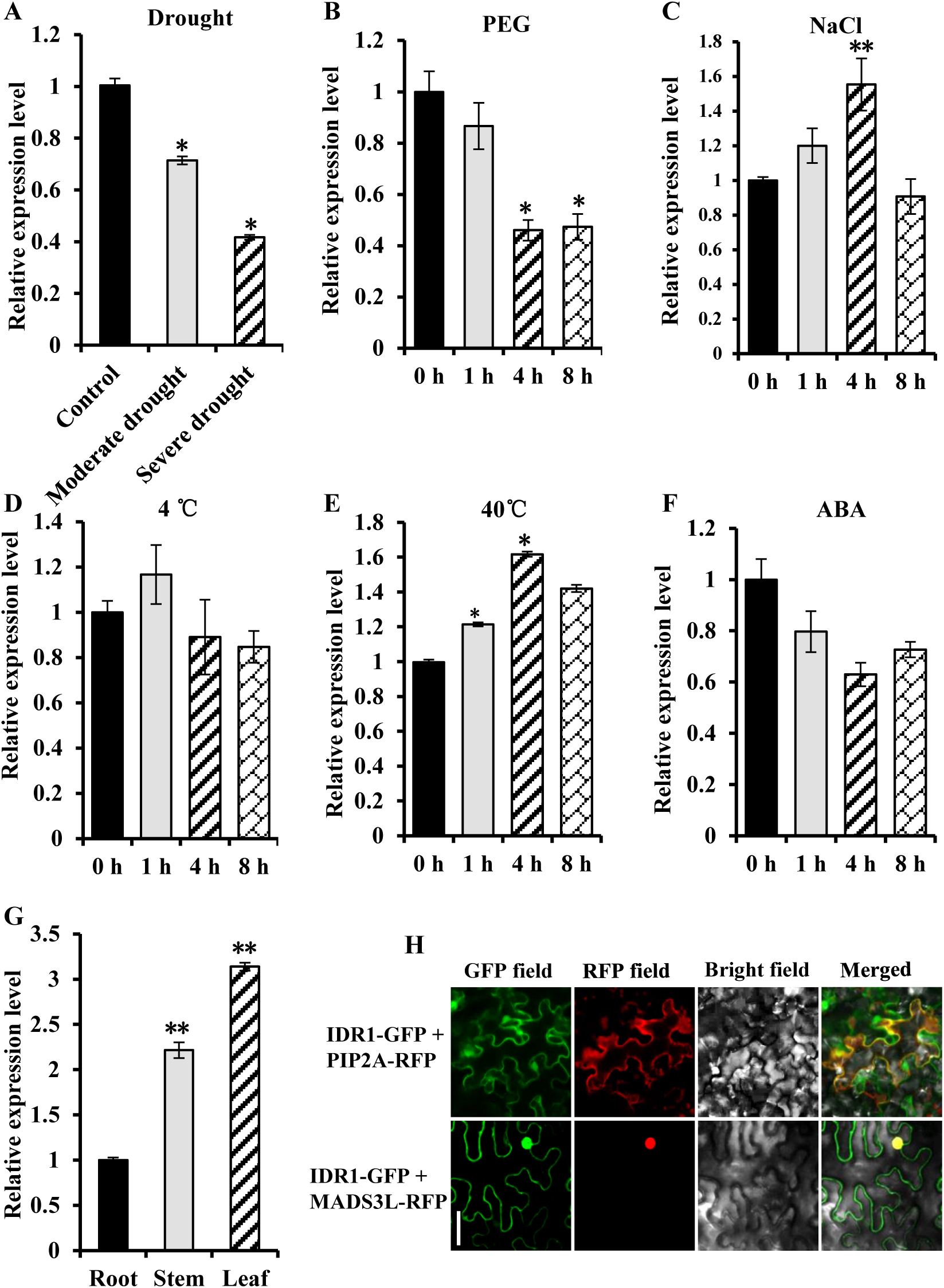
Expression patterns and subcellular localization of IDR1. (A-F) Expression patterns of *IDR1* after treatment with drought, PEG, NaCl, 4°C, 40°C or ABA. Leaves were sampled from 2-week-old IAPAR9 seedlings treated with moderate or severe drought stress (A), 20% PEG (B), 150 mM NaCl (C), 4°C (D), 40°C (E) or 50 μM ABA (F) at the indicated time points (0, 1, 4, 8 h) following the treatments, and then total RNA was isolated and subjected to qRT-PCR assays. Control, no drought stress. **(G)** Tissue-specific expression patterns of *IDR1* at seedling stage. Total RNA extracted from roots, stems and leaves of 4-week-old seedlings was used for qRT-PCR assays. **(H)** Subcellular localization of IDR1. *IDR1* cDNA was fused to *GFP*, and the resulting *IDR1-GFP* construct was co-transfected with *PIP2A-RFP* (plasma membrane localization marker) (upper panels) or *MADS3L-RFP* (nuclear localization marker) (lower panels) into 30-d-old *N. benthamiana* leaves for fluorescence observation. PIP2A, plasma membrane intrinsic protein 2A; MADS3L, MADS-box transcription factor 3-like protein. Scale bar = 20 μm. Data are means (±SD) for three biological replicates. Significant differences from control (A), 0 h (B-F) or root (G) were determined by Student’s *t*-test: *P < 0.05, **P < 0.01, t-test.

In order to get a clearer picture of expression patterns of IDR1, tissue-specific expression patterns of *IDR1* were examined by qRT-PCR assays. The results demonstrated that *IDR1* was ubiquitously expressed in all the three tested tissues, with the highest expression in leaves and the lowest expression in roots (Fig. 3G). Assays of subcellular localization of IDR1 revealed that IDR1 was definitely localized in nucleus, and also at plasma membrane or cell periphery (Fig. 3H). In addition, it seems likely that IDR1 was localized in cytoplasm as well (Fig. 3H). These observations suggest that IDR1 presumably performs particular functions on plasma membrane and nucleus.

### *idr1-1* Mutation Results in Physiological and Morphological Changes that are Associated with Drought Tolerance

To learn about physiological changes caused by *idr1-1* mutation, we determined the water potentials of the genotypes as indicated in Fig. 4A. Under control conditions, leaves of *idr1-1* mutants had relatively higher water potential values in comparison with those of IAPAR9 plants and the complementation lines at 10 o’clock onwards (Fig. 4A, left panel); a similar trend was observed when under moderate drought conditions: the *idr1-1* mutant leaves possessed quite obviously higher water potential values between 10 to 18 o’clock (Fig. 4A, right panel), which collectively indicates that the *idr1-1* mutant leaves has greater capability to retain water than do leaves of the IAPAR9 plants and the complementation lines. Besides, water loss rates of detached leaves from *idr1-1* mutants were markedly lower than those from IAPAR9 plants and the three complementation lines, from 2 h onwards after sampling; at time point 12 h, the water loss rate was approximately 72% for IAPAR9 leaves, as against about 60% for *idr1-1* leaves (Fig. 4B), suggesting that the increased drought tolerance of the *idr1-1* mutant may be, at least in part, a result of attenuated water loss rates of leaves. Although proline contents exhibited no differences among these genotypes under control conditions, the content in *idr1-1* leaves was significantly higher than the contents in the leaves of other 4 genotypes under the moderate drought conditions (Fig. 4C), indicating that drought stress might have promoted biosynthesis of proline in the *idr1-1* mutant background. Further, we measured malonaldehyde (MDA) contents as well as relative electrolyte leakage (REL) of leaves, and found that *idr1-1* mutants possessed a significantly lower MDA level and REL compared to the remaining genotypes under the moderate drought conditions (Fig. 4, D and E). In addition, *idr1-1* mutants, which constantly possessed dark-green leaves, were of significantly higher chlorophyll contents compared with the other genotypes when grown in both control and moderate drought conditions (Fig. 4F). All these data together demonstrate that *idr1-1* mutants exhibit significantly enhanced capability of retaining water and preserving integrity of plasma membrane, which is presumably one of the important reasons why *idr1-1* mutants show elevated drought tolerance.

**Figure 4.**
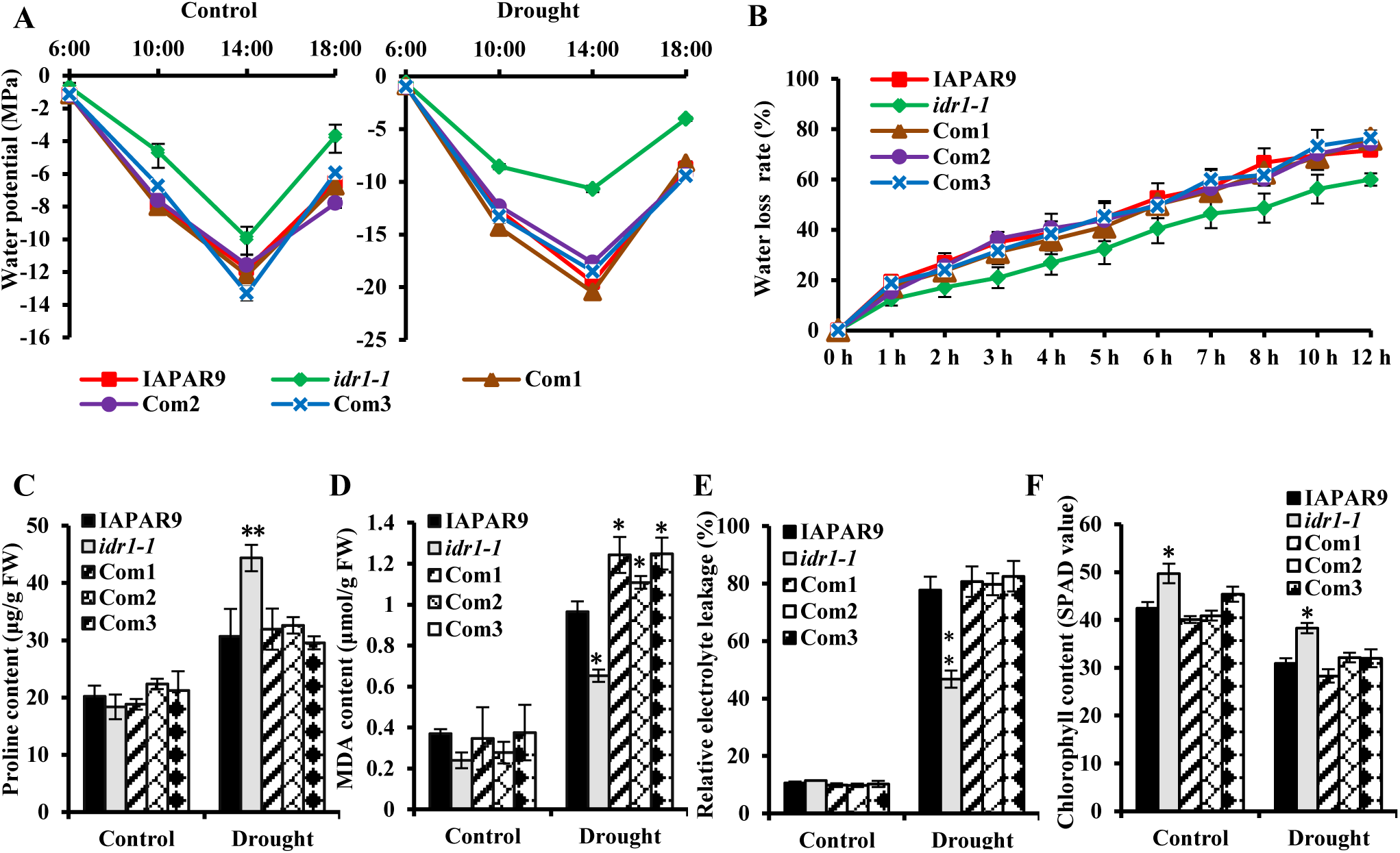
Physiological changes of *idr1-1* mutant help retain more water in its leaves. **(A)** Measurements of leaf water potentials of *idr1-1* mutants, IAPAR9 plants, and *IDR1* complementation lines grown under control or moderate drought stress conditions. Leaf samples for measuring water potentials were collected at the 4 time points as indicated. Control, no drought stress. **(B)** Measurements of water loss rates of detached leaves from *idr1-1* mutants, IAPAR9 plants, and *IDR1* complementation lines at the indicated time points. Water loss was expressed as the percentage of decreases in fresh weights (FW) of the detached leaves at the different time points compared to their own at 0 h. Data are means (±SD) for three biological replicates; for each genotype, 6 leaves were used for measuring leaf water potential (A) or leaf FW (B) in each of the three replicates. **(C-F)** Measurements of proline contents (C), MDA contents (D), REL (E) and relative chlorophyll contents (F) in leaves of *idr1-1* mutants, IAPAR9 plants and *IDR1* complementation lines under control or moderate drought stress conditions. Data are means (±SD) for three biological replicates. Significant differences from IAPAR9 in each group (control or drought) of every bar chart were determined by Student’s *t*-test: *P < 0.05, **P < 0.01, t-test. Control, no drought stress.

Under control conditions, it appeared that stomatal apertures of *idr1-1* leaves showed a slight, but not significant, reduction compared to the other genotypes as indicated (Fig. 5, A and B); however, they were significantly greater than those of the other genotypes under drought stress conditions (Fig. 5, A and B, upper panel), which may stems from an impairment of signaling of stomatal closure. There were no evident differences in stomatal densities between *idr1-1* mutant and the rest of genotypes under control or drought conditions (Fig. 5B, lower panel). Taken together, these results suggest that mutation of *IDR1* leads to attenuated stomatal closure, but does not significantly alter the stomatal density.

**Figure 5.**
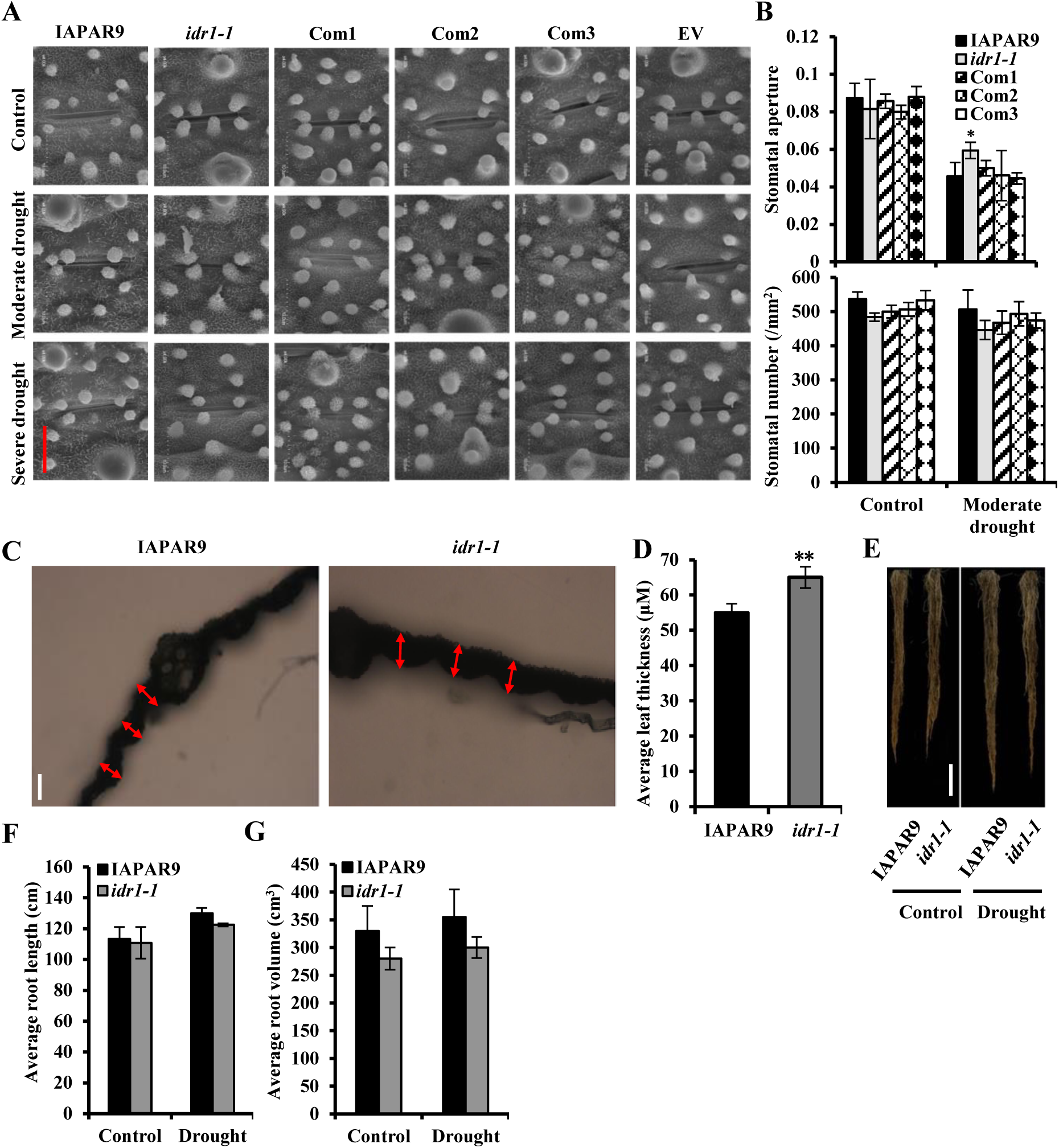
Morphological changes contribute to enhanced drought tolerance in *idr1-1* mutant. **(A)** Representative images for stomatal apertures of leaves of *idr1-1* mutants, IAPAR9 plants and *IDR1* complementation lines under control or drought conditions. The photographs were taken with a scanning electron microscope. Scale bar = 10 μm. Control, no drought stress. **(B)** Statistics of stomatal apertures (upper panel) and stomatal numbers (lower panel) of *idr1-1* mutants, IAPAR9 plants and *IDR1* complementation lines, which were derived from more stomatal images taken simultaneously with those in A. Data are means (±SD) for three biological replicates; for each genotype, 6 stomata were used for measuring stomatal apertures in each of the three replicates. **(C and D)** Average thickness of transverse sections of *idr1-1* mutant and IAPAR9 leaves. The thickness of *idr1-1* mutant and IAPAR9 flag leaves at the positions in the proximity of upper portions of the main vein were measured (C) and shown (D). Scale Bar = 50 μm. **(E-G)** Effects of *idr1-1* mutation on root traits. Plants were grown in PVC tubes for 1 month, and then they were treated with or without (control) moderate drought stress for another 2-3 weeks by withholding water. The whole roots were isolated from soils by washing them with running water, and then imaged (E). Root lengths and root volumes of *idr1-1* mutant and IAPAR9 plants were measured and the means of them were shown (F and G). Data are means (±SD) for three biological replicates; for each genotype, 5 leaves (D) or 3 plants (F and G) were used for measuring leaf thickness or root lengths as well as root volumes, respectively, in each of the three replicates. Significant differences from IAPAR9 were determined by Student’s *t*-test: **P < 0.01, t-test. Scale Bar = 20 cm.

It is widely known that thicker leaves have beneficial effects on plant tolerance to drought stress as they are able to retain more water to provide protection against soil water deficiency. As depicted in Fig. 5, C and D, *idr1-1* leaves looked thicker than IAPAR9 leaves at the position in the proximity of upper portion of the main vein; average thickness of *idr1-1* leaves reached 65 μm, being 10 μm longer than that of IAPAR9 leaves (Fig. 5D), which may be one of the reasons that *idr1-1* mutants exhibits strongly enhanced drought tolerance relative to IAPAR9 plants. Besides, deposition of wax crystals on the front and back of *idr1-1* leaves was slightly, but not significantly, less than that of IAPAR9 leaves, implicating that mutation of *IDR1* has little effects on the deposition of wax crystals on leaves (Supplemental Fig. S6). To know the differences in roots between *idr1-1* mutants and IAPAR9 plants, we determined the several traits of roots that were derived from PVC tube-grown *idr1-1* mutants and IAPAR9 plants (Supplemental Fig. S7A). Average root lengths of both *idr1-1* mutants and IAPAR9 plants grown in moderate drought conditions were slightly increased compared to that of both genotypes grown in control conditions (Fig. 5, E and F); however, *idr1-1* mutants and IAPAR9 plants showed no significant differences in the average root length when both genotypes were grown in either control or drought conditions (Fig. 5, E and F). Although *idr1-1* mutants showed a slight decrease in average root volume and a significant decrease in average root number relative to IAPAR9 plants, moderate drought treatments seemingly had only slight or no significant impacts on root volumes or root numbers of both genotypes, respectively (Fig. 5G; Supplemental Fig. S7B); as shown in Fig. 5G, the root volumes of both *idr1-1* mutants and IAPAR9 plants undergoing the drought stress were a little larger than those of both the genotypes grown in control conditions. Furthermore, the drought treatments also induced slight increases in root dry weights and root diameters of *idr1-1* mutants and IAPAR9 plants in comparison with non-drought treatments (control) (Supplemental Fig. S7, C and D); it appeared that mutation of *IDR1* led to small increases in root diameters, but subtle decreases in root dry weights under both control and drought conditions (Supplemental Fig. S7, C and D). Considering that *idr1-1* mutants have a little changed average root length, average root volume, average root dry weight and average root diameter, together with thicker leaves and remarkably reduced plant heights, the mutant plants must have less transpirational water loss but comparable water absorption from soil when compared to IAPAR9 plants, which therefore makes *idr1-1* more tolerant of drought stress.

### *idr1-1* Mutation Impairs Apoplastic and Chloroplatic ROS Production Provoked by Drought Stress and Oxidative Stress Inducer

To gain insights into the status of ROS accumulation and cell death in *idr1-1* mutants when challenged by drought stress, we performed DAB, NBT, and Trypan blue staining of the leaves from different genotypes [treated with or without (control) drought stress] as indicated in Fig. 6A. The results from DAB staining demonstrated that the leaves from control did not show visible H_2_O_2_ accumulation as they lacked staining, but moderate drought treatments provoked a stronger accumulation of H_2_O_2_ in the leaves of IAPAR9 plants, complementation lines and the OE lines when compared with the *idr1-1* leaf (sample 2) that was less stained (Fig. 6A, left panel). NBT staining revealed that O_2_^-^ production in *idr1-1* leaf was less than in leaves of the remaining genotypes after drought treatments (Fig. 6A, middle panel). A similar trend was observed when the leaves from these genotypes were stained with Trypan blue to detect the occurrence of cell death under control and drought stress conditions, that is to say, *idr1-1* leaf accumulated less amounts of Trypan blue stain compared with the rest of these genotypes under drought stress conditions (Fig. 6A, right panel). These data collectively suggest that *idr1-1* mutation attenuates accumulation of the major species (H_2_O_2_ and O_2_^-^) of leaf ROS induced by drought stress, and such mutation mitigates cell death elicited by the stress.

**Figure 6.**
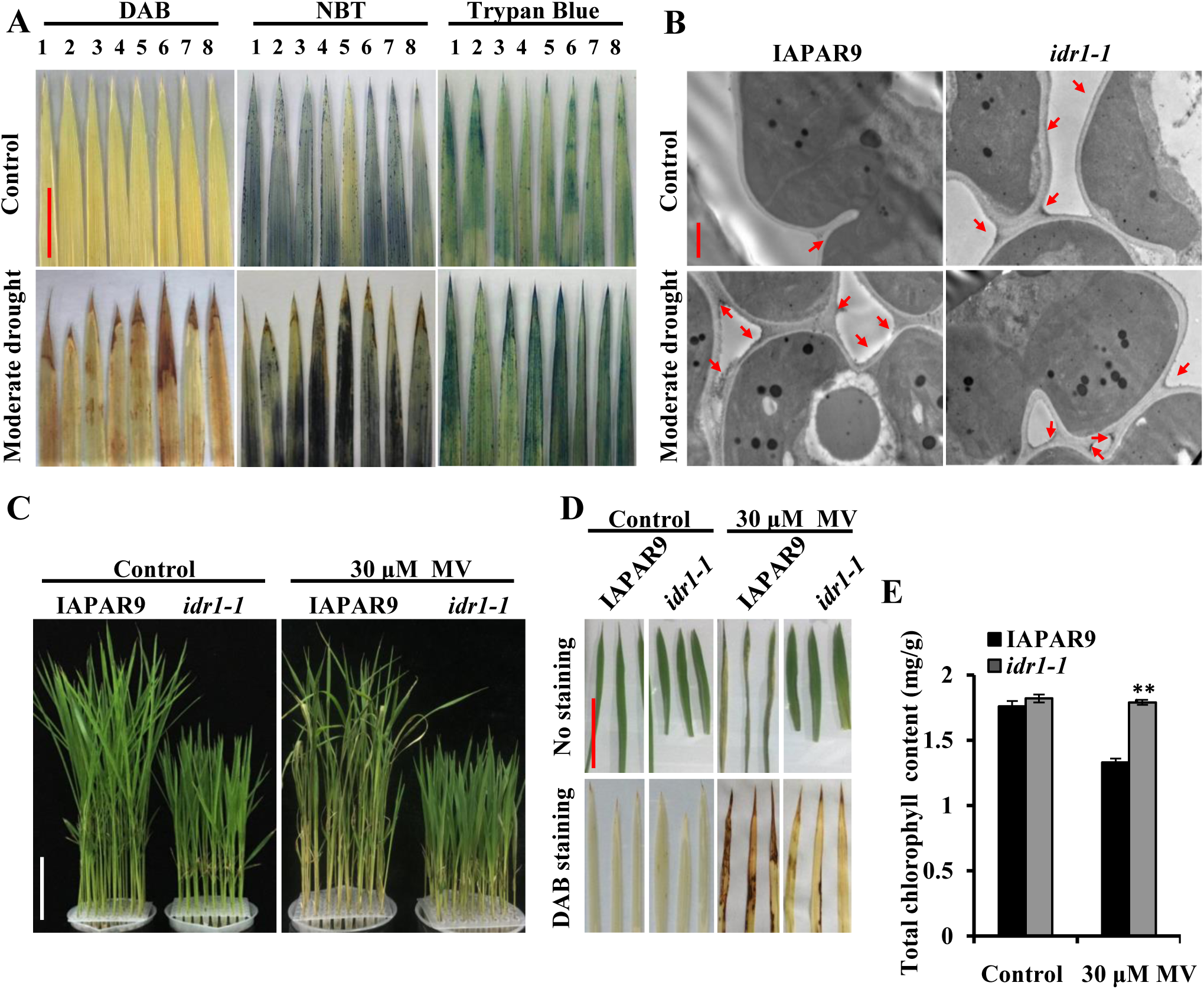
*idr1-1* mutation attenuates apoplastic and chloroplastic ROS production triggered by drought stress and MV. **(A)** DAB, NBT and Trypan blue staining of detached leaves from *idr1-1* mutant and IAPAR9 plants. Five-week-old *idr1-1* mutant and IAPAR9 seedlings underwent drought stress for about 2 weeks (moderate stress) and the detached leaves of IAPAR9, *idr1-1*, Com1, Com2, Com3, OE2, OE3 and OE5 (1-8, respectively) were used for such staining. Scale Bar = 10 cm. **(B)** Cytochemical localization of drought-induced H_2_O_2_ in intercellular space between mesophyll cells of *idr1-1* mutant or IAPAR9 leaves. Leaf samples for sectioning were from the experiment depicted in A. The H_2_O_2_ was stained with CeCl_3_ and the photographs were taken with transmission electron microscopy (TEM). Red arrows indicate CeCl_3_ precipitates. Scale Bar = 1 μm. **(C-E)** Performance of *idr1-1* mutant and IAPAR9 seedlings after MV treatments. Two-week-old *idr1-1* mutant and IAPAR9 seedlings cultured in liquid Hoagland medium were treated with 30 μM MV for 3 days (C). DAB staining of leaves from *idr1-1* mutant and IAPAR9 seedlings following MV treatments were performed to detect amounts of H_2_O_2_ (D). In parallel, total chlorophyll contents of the *idr1-1* mutant and IAPAR9 leaves from the experiment shown in C were measured and shown (E). Data are means (±SD) for three biological replicates; for each genotype, 30 plants were used for measuring leaf chlorophyll contents in each of the three replicates. Significant differences from IAPAR9 were determined by Student’s *t*-test: **P < 0.01, t-test. Scale Bars = 10 cm (C and D).

We further examined subcellular localization of H_2_O_2_ accumulation in the leaves undergoing moderate drought treatments by means of a cytochemical technique. It is apparent from the Fig. 6B that, under control (unstressed) conditions, leaf apoplasts of IAPAR9 plants did not accumulate visible CeCl3 deposits, while those of *idr1-1* mutants did indeed accumulate a certain number of CeCl3 deposits, indicative of higher H_2_O_2_ accumulation in the *idr1-1* leaf apoplasts (Zhang et al., 2010). This suggest that mutation of *IDR1* might have evoked biosynthesis of a particular amount of H_2_O_2_ in leaf apoplasts without drought stress, which may provide a reasonable explanation for the above observation that stomatal apertures of *idr1-1* mutant leaves slightly smaller than those of IAPAR9 leaves under control conditions (Fig. 6B). By contrast, under drought stress conditions, a relatively larger number of CeCl3 deposits were observed in the leaf apoplasts of IAPAR9 plants than of *idr1-1* mutants, suggesting that more H_2_O_2_ in the leaf apoplasts are accumulated in response to drought stress in the IAPAR9 leaves. It should be noted that the accumulation of CeCl3 deposits showed no significant differences between the *idr1-1* samples with or without the treatment of drought stress, suggesting that drought stress does not trigger the H_2_O_2_ accumulation efficiently in the *idr1-1* leaves, which is in agreement with the results of DAB staining (Fig. 6B).

After 2-week-old *idr1-1* mutant and IAPAR9 seedlings were treated with or without (control) 30 μM methyl viologen (MV) for 72 h, the seedlings of IAPAR9 became noticeably wilted and/or yellowing following treatments, while those of *idr1-1* mutants were affected to a lesser extent (Fig. 6C). DAB staining revealed that detached leaves from IAPAR9 plants experiencing MV treatments showed much greater ROS production than those from *idr1-1* mutants, which was marked by a much larger area of intense staining existing on IAPAR9 leaves than on *idr1-1* leaves (Fig. 6D). In addition, the IAPAR9 leaves subjected to MV treatment possessed a noticeably lower chlorophyll content than those without MV treatment; however, there were almost no differences in chlorophyll contents between MV-treated and untreated *idr1-1* leaves, indicating that chlorophyll in IAPAR9 leaves is obviously broken down after treatment with MV (Fig. 6E). Given the fact that MV is able to elicit rapid burst of chloroplastic H_2_O_2_ production by inhibiting photosynthetic electron transport (PET), the results mentioned above thus suggest that *idr1-1* mutation markedly attenuates MV-induced H_2_O_2_ generation in chloroplasts, thus diminishing oxidative damage to leaf tissues and chloroplasts themselves.

When *idr1-1* mutant seeds developed to full maturity (husks turned fully yellow) in the upland fields, *idr1-1* mutant leaves remained green and became hardly senescent, while the IAPAR9 control leaves had become almost totally yellow as a result of senescence at the same developmental stage, indicating that *idr1-1* mutants have much delayed leaf senescence and breakdown of proteins as well as membranous structures following the seed maturity. Moreover, upon 3-d-darkness induction, IAPAR9 leaves displayed greater leaf senescence than did *idr1-1* leaves (Supplemental Fig. S8, top panel). DAB staining revealed that *idr1-1* leaves lacked staining compared with strongly stained IAPAR9 leaves (Supplemental Fig. S8, middle panel); NBT staining also showed that *idr1-1* leaves were stained less than IAPAR9 leaves (Supplemental Fig. S8, bottom panel). Altogether, these data suggest that the delayed leaf senescence induced by darkness may result from lack or attenuation of ROS accumulation in *idr1-1* mutants, which is in accord with the notion that *idr1-1* mutation impairs ROS production and/or increases ROS-scavenging ability.

A recent genome-wide analysis revealed that there existed ten and nine respiratory burst oxidase homologs (RBOHs) in Arabidopsis and rice, respectively. All the 9 rice *OsRBOHs* (*OsRBOHA-OsRBOHI*) were differentially expressed during the entire life cycle of rice (Chang et al., 2016). The expression of five of the rice *OsRBOHs*, i.e. *A-C*, *E* and *I*, were induced to higher levels by drought stress, with expression levels under drought stress being about 10, 4, and 8.5 times as high as those under normal conditions, respectively (Wang et al., 2013). Our data, however, showed that among the five tested genes, only the *OsRBOHB* in *idr1-1* mutants was induced to a level approximately twice as high as that in IAPAR9 plants under severe drought stress conditions (Supplemental Fig. S9; Supplemental Table S2). Thus, it appears that drought stress fails to induce expression of *OsRBOHs* efficiently in the *idr1-1* mutant background, which may be one of the reasons why *idr1-1* mutant produces less amounts of ROS upon exposure to drought stress.

Chloroplast is known as one of the major sources of ROS. During photosynthesis, especially when plants are subjected to abiotic stress, ROS are generated in abundance as a result of impaired electron transport (Challabathula et al., 2018). To know if the enhanced drought tolerance of *idr1-1* mutant is associated with altered photosynthetic process, we examined expression levels of 11 genes that were involved in chloroplastic PET or redox reactions. Nine of the eleven tested genes, i.e. *OsLFNR1*, *OsFd4*, *OsNTRC*, *OsPC*, *OsGLT2*, *OsCDSP32*, *OsTrxX*, *OsTrxM5* and *OsABC1*, showed higher expression levels in *idr1-1* mutants than in IAPAR9 plants under moderate and severe drought stress conditions (Supplemental Fig. S10; Supplemental Table S2). Among the 9 genes, five genes (*OsLFNR1*, *OsPC*, *OsGLT2*, *OsTrxM5* and *OsABC1*) were expressed at least 2 times as high in *idr1-1* mutants as they were in IAPAR9 plants under moderate and/or severe drought stress conditions (Supplemental Fig. S10). These results together suggest that under drought stress conditions, the expression of these 9 photosynthesis-related genes in *idr1-1* mutants was induced to higher levels when compared to the levels in IAPAR9 plants. *OsLFNR1* encodes a chloroplast-localized leaf-type ferredoxin-NADP^+^-oxidoreductase, and it is able to transfer an electron from ferredoxin (Fd) molecules to NADPH for use in Calvin-Benson cycle. *OsNTRC* encodes a bifunctional thioredoxin reductase that is required for reduction of oxidized thioredoxins. *OsFd* encodes a 2Fe-2S iron-sulfur cluster binding domain containing protein, and it receives electrons from PSI and then transfers them to OsFNR (or OsLFNR) or to O_2_ to form O_2_^-^. *OsPC* encodes a plastocyanin, which transfers electrons from Cytb6f complex to PSI. The remainder are thioredoxin- or thioredoxin-related genes that function in Calvin-Benson cycle or H_2_O_2_-detoxifying system (Gong et al., 2020). It should be noted that OsFd can transfer electrons from PSI to OsLFNR in normal circumstances, or to O_2_ to produce excessive O_2_^-^ under stress conditions due to inhibition of Calvin-Benson cycle. Thus, it appears that the elevated expression levels of the genes (e.g. *OsLFNR1*, *OsFd4* and *OsPC*) may contribute to enhanced PET for successful generating NADPH and ATP for Calvin-Benson cycle, and thereby diminish the transfer of electrons to O_2_ for eventual H_2_O_2_ overproduction. Moreover, the upregulation of thioredoxin- or thioredoxin-related genes may continuously maintain Calvin-Benson cycle in an active state and limit the overaccumulation of H_2_O_2_ against undue oxidative stress on chloroplasts.

### Increased Expression of the Genes Associated with ROS Scavenging in *idr1-1* Mutant Diminishes ROS Accumulation Caused by Drought Stress

In order to understand the molecular mechanisms underlying enhanced drought tolerance observed in *idr1-1* mutant, we conducted gene expression analysis to examine the levels of those genes associated with ROS scavenging and PET. Three-week-old seedlings were treated with moderate or severe drought stress and subsequently used for qRT-PCR assays. The results demonstrated that two marker genes, *RAB16C* and *RAB16A*, were highly induced by severe drought treatment, indicating that drought treatment is quite effective (Supplemental Fig. S11; Supplemental Table S2); another three genes tested, *OsLEA3-1*, *OsLG3*, and *DSM1*, were all induced to higher expression levels in *idr1-1* mutants than in IAPAR9 plants under severe drought stress conditions (Fig. 7A; Supplemental Table S2). OsLG3 was discovered to positively regulate drought tolerance in rice by inducing gene expression of a dozen ROS scavengers (Xiong et al., 2018). DSM1 was required for the normal expression of two ROS-scavenging genes, *OsPOX22.3* and *OsPOX8.1* (Ning et al., 2010). LEA3-1 was considered as a protectant for desiccation and oxidation stress (Jangam et al., 2016). To see whether *idr1-1* mutant has increased expression levels of the ROS-scavenging genes due to mutation of *IDR1*, we further checked the transcript levels of a dozen or so genes implicated in ROS scavenging. It is quite interesting that 11 genes, including 7 APX-family genes (*OsAPX1-OsAPX2, OsAPX4-OsAPX8*), two CAT-family genes (*OsCAT1* and *OsCAT3*), one *OsFeSOD* as well as one *OsPOX22.3* genes, showed higher expression levels in *idr1-1* mutants than in IAPAR9 plants under moderate and/or severe drought stress conditions (Fig. 7B; Supplemental Table S2). *APX* genes encode ascorbate peroxidases that are able to use ascorbate to reduce H_2_O_2_ to H2O; *CAT* genes encode catalases that are highly efficient at catalyzing breakdown of H_2_O_2_ to H_2_O and O_2_; FeSOD is very active in converting O_2_^-^ into H_2_O_2_, and POX actively acts on H_2_O_2_ to make it harmless H_2_O via redox reactions (Ning et al., 2010). Furthermore, measurements of POD enzymatic activity revealed that under either control or moderate drought conditions, POD activities in *idr1-1* mutants were always significantly higher than in IAPAR9 plants (Fig. 7C). Altogether, these data provide a clear demonstration that mutation of *IDR1* might have led to activation of ROS-scavenging system, and thereby induces elevated expression of such important genes for highly efficient ROS scavenging and rise in POD activity, which is presumably one of the important causes of the significantly enhanced tolerance of *idr1-1* mutant to drought stress.

**Figure 7.**
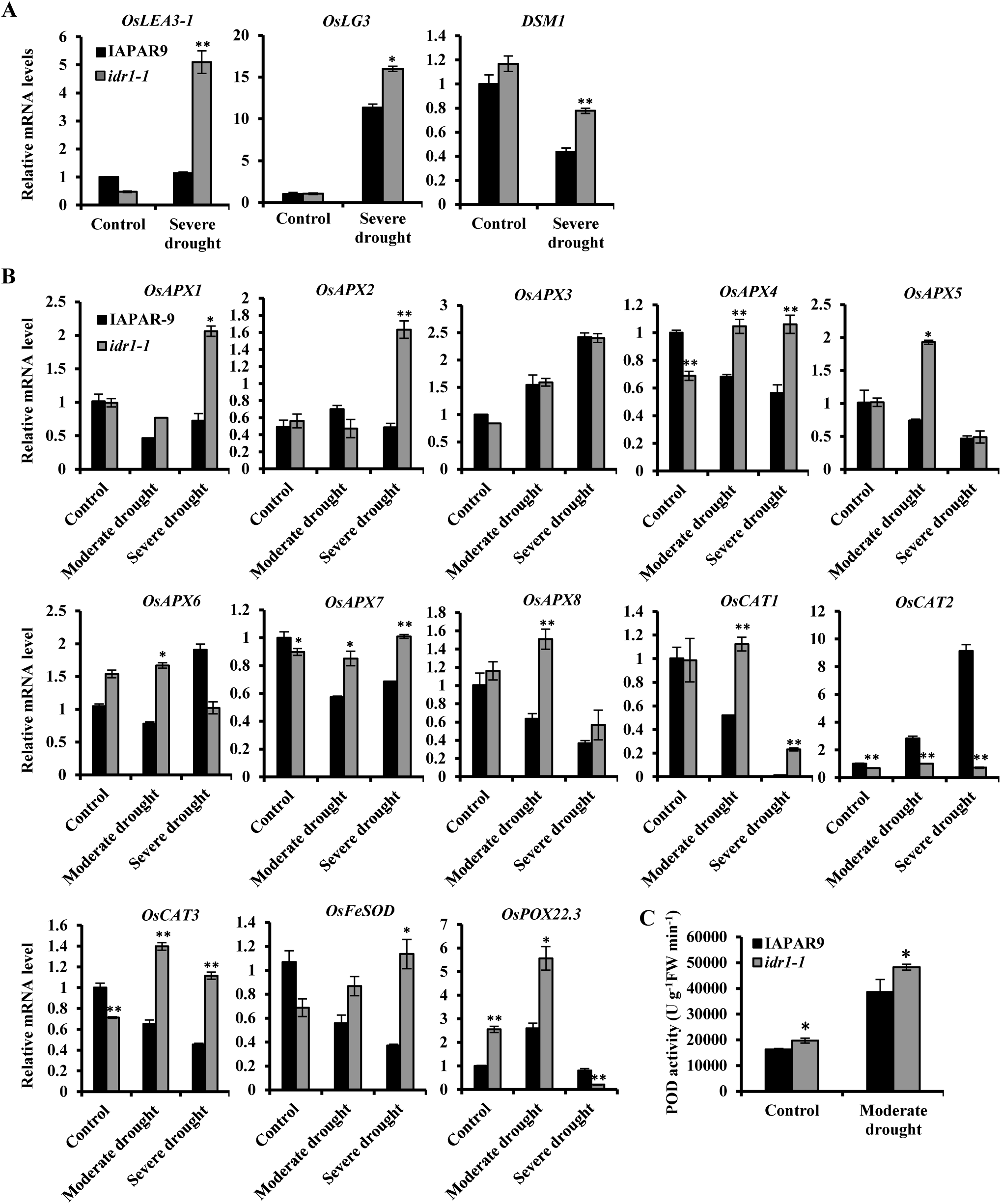
*idr1-1* mutation increases expression of a group of ROS-scavenging genes under drought-stressed conditions. **(A)** Relative expression levels of three genes that were reportedly associated with activation of ROS-scavenging system. **(B)** Relative expression levels of a dozen genes involved in ROS scavenging under control or drought stress (moderate or severe) conditions. For qRT-PCR analyses in A and B, leaves from 4-week-old seedlings of *idr1-1* and IAPAR9 experiencing nonstressed (control) and drought-stressed treatments, were used for RNA isolation and qRT-PCR assays to examine expression of the genes as indicated. Rice *ubiquitin 5* gene was amplified and used as an internal control. **(C)** POD activity assays in *idr1-1* mutant and IAPAR9 leaves under control or moderate drought stress conditions. Leaf samples for POD activity assays were from the experiment described in B. Data are means (±SD) for three biological replicates. Significant differences from IAPAR9 were determined by Student’s *t*-test: *P < 0.05, **P < 0.01, t-test.

### Transcriptome Analyses of Drought-stressed *idr1-1* Mutants Reveals Several Mechanisms Underlying Enhanced Drought Tolerance

To further understand the underlying mechanism for the enhanced drought tolerance caused by *idr1-1* mutation, we examined transcriptome profiles of 5-week-old *idr1-1* mutants and IAPAR9 plants that had undergone 14-d progressive drought stress treatment by using whole-genome mRNA sequencing (mRNA-seq) method (Supplemental Fig. S12). The analyses of the transcriptomic data demonstrated that there were 941 and 120 differential expressed genes (DEGs) identified under control and drought-stressed conditions, respectively, which separately generated a [Control *idr1-1* vs Control IAPAR9] (Supplemental Data Set S1) and a [Drought *idr1-1* vs Drought IAPAR9] data sets (Supplemental Data Set S2). Further analyses revealed that a total of 397 upregulated and 544 downregulated genes were found in the [Control *idr1-1* vs Control IAPAR9] data set (Supplemental Data Set S1); likewise, there were 45 upgreulated and 75 downregulated genes present in the [Drought *idr1-1* vs Drought IAPAR9] data set (Supplemental Data Set S2). This suggests that *idr1-1* mutation largely leads to downregulation of a group of genes in the *idr1-1* mutant in comparison with IAPAR9 control. We then conducted Gene Ontology (GO) analyses to classify these DEGs. The results demonstrated that the 941 DEGs from the [Control *idr1-1* vs Control IAPAR9] data set were significantly enriched for oxygen binding, secondary metabolic process, and cell growth GO terms (Supplemental Fig. S13); the 120 DEGs from the [Drought *idr1-1* vs Drought IAPAR9] data set were significantly enriched for secondary metabolic process (Supplemental Fig. S13). Under drought-stressed conditions, 11 upregulated genes and 38 downregulated genes, the expression of which were affected by both *idr1-1* mutation and drought stress, were obtained by overlapping the DEGs from the three different data sets as indicated (Supplemental Fig. S13; Supplemental Table S3). To ascertain how transcriptional changes of a collection of genes in *idr1-1* mutant increased its drought tolerance, we carefully examined the genes in both [Control *idr1-1* vs Control IAPAR9] and [Drought *idr1-1* vs Drought IAPAR9] data sets, and discovered a total of 43 genes that were presumably involved in drought tolerance (Supplemental Table S4). Although some genes in Supplemental Table S4 only showed significantly differential expression in control conditions but not in drought conditions, the altered expression of such genes, however, will help *idr1-1* mutants to cope better with drought stress, at least in part, at the onset of occurrence of drought stress due to greater capacity of ROS scavenging and antioxidation in the *idr1-1* mutants. These genes were then grouped into 7 classes on the basis of reported or putative functions about them: (1) reduction of oxidative stress, ROS production, cell death, and hypersensitive reaction (Class I); (2) increase of ROS scavenging and detoxification (Class II); (3) increase of antioxidant biosynthesis (flavonoid, anthocyanin, etc.) (Class III); (4) enhancement of osmotic adjustment (Class IV); (5) weakening of leaf senescence (Class V); (6) alteration of hormonal biosynthesis, signaling and response (Class VI); and (7) elevation of water retaining, survival ability, and drought resistance (Class VII) (Supplemental Table S4)..

There are 11 genes probably associated with the functions shown in Class I (Supplemental Table S4). Six genes, which encode OsOXO4 (Os03g48780), OsDGK8 (Os12g12260), OsCBSX10 (Os01g44250), UDP-glucosyltransferase (Os02g11680), Yr10-like protein (Os11g37774) and OsRING-1 (Os02g52210), respectively, showed decreased expression in drought-stressed conditions (Supplemental Table S4). OsOXO4 was reported to contribute to apoplastic ROS accumulation under drought conditions (Voothuluru and Sharp, 2013; Zhang et al., 2013). OsDGK8 plays a role in generating phosphatidic acid (PA) in response to biotic and abiotic stresses (Li et al., 2015), suggesting that it may participate in PA-dependent ROS production. OsCBSX10, UDP-glycosyltransferase, Yr10-like protein, OsRING-1 and their Arabidopsis homologs were documented to perform functions in mitochondrial ROS accumulation, pathogen-induced oxidative burst as well as hypersensitive responses (Meng et al., 2006; Coram et al., 2010; Shin et al., 2020). Thus, these data collectively suggest that downregulation of such genes in *idr1-1* mutant presumably attenuates ROS production and cell death in different cellular compartments or tissues. Four genes, which encode OsPLC4 (Os05g03610), OsAAO3 (Os07g18154), NBS-LRR disease resistance protein (Os06g49390) and NB-ARC domain containing protein (Os11g34970), were all downregulated in *idr1-1* mutant only under control conditions (Supplemental Table S4). OsPLC4 is a phospholipase C, and it was reportedly associated with ROS production upon infection by bacterial strain *Pst* DC3000 (D’Ambrosio et al., 2017). OsAAO3 is an aldehyde oxidase 3, and it can oxidize NADH, ultimately leading to generation of O_2_^-^ (Kundu et al., 2012; Batth et al., 2017). Os06g49390 and Os11g34970 both encode disease resistance proteins; overexpression of this class of genes brings about rapid HR, cell death and growth retardation (Bai et al., 2012). Hence, reduced expression of the above-mentioned 4 genes in *idr1-1* mutant is helpful in weakening ROS generation and cell death. Os01g64120 encodes a 2Fe-2S iron-sulfur cluster binding domain containing protein (Fd-like protein) for facilitating electron transfer from PSI to FNR in normal circumstances, and upregulated expression of such genes in *idr1-1* mutant under both drought and control conditions may prevent overproduction of O_2_^-^ (Hanke and Mulo, 2013). Taken together, these results suggest that altered expression of the foregoing genes in *idr1-1* mutant may be favorable to lessening ROS production and cell death under drought stress conditions. There are 10 genes (Class II), which have putative functions in ROS scavenging or detoxification, showing altered expression under drought-stressed or control conditions (Supplemental Table S4). Nine genes, i.e. *OsPOX71* (Os05g04500), *OsANN3* (Os05g31750), *OsPOX62* (Os04g59200), *OsPOX8.1* (Os07g48010), *OsPOX22.3* (Os07g48020), *OsPOX3006* (Os07g48050), *OsGSTU6* (Os01g37750), *OsLG3* (Os03g08470), and *OsMT2d* (Os01g05585) displayed upregulated transcription in *idr1-1* mutant under drought or control conditions. Peroxidases (including OsPOX71, OsPOX62, OsPOX8.1, OsPOX22.3 and OsPOX3006) fulfilled the functions of catalyzing conversion of H_2_O_2_ to H_2_O (Atack and Kelly, 2007; Hasanuzzaman et al., 2017; Majumdar and Kar, 2019) (Supplemental Table S4). Overexpression of *OsANN3* in rice led to elevated activities of ROS-scavenging enzymes, including APXs, CATs and SODs (Li et al., 2019). OsGSTU6 was involved in scavenging H_2_O_2_ and reducing oxidation stress (Ning et al., 2010). OsLG3 was recently reported to contribute positively to upregulation of 10 of ROS scavenging-related genes (like *APXs*, *CATs*, *PODs*, *SODs*, etc.) upon overexpression of *OsLG3* (Xiong et al., 2018). OsMT2d performed a function in decreasing oxidative damage when rice plants were exposed to abiotic stress conditions (Patankar et al., 2019). *OsAAO1* (Os06g37080), which showed decreased expression in control conditions, encodes an ascorbate oxidase 1. This enzyme catalyzes active L-ascorbate to dehydroascorbate, and the accumulation of such dehydroascorbates was correlated with leaf senescence and H_2_O_2_ elevation (Chen et al., 2013). Altogether, these data demonstrate that expression changes of the foregoing genes in *idr1-1* mutant appear to make positive contributions to *idr1-1*’s tolerance against drought stress.

Antioxidants play important roles in reducing oxidative damage and eliminating free radicals when plants are subjected to abiotic stress. Our analyses led to the discovery of 5 genes (Class III) associated with antioxidant biosynthesis or production, which were all upregulated in drought or control conditions (Supplemental Table S4). Four genes, Os07g02100 and *OsF3H2* (Os10g39140) (upregulated in drought conditions), as well as Os01g13610 and Os06g44170 (upregulated in control conditions), are all involved in flavonoid biosynthesis (Supplemental Table S4). Flavonoids played important roles in protecting cell wall and membranes when plants underwent stresses (Song et al., 2016). OsF3H2 in woody plant *Lycium chinense* had strong effects on scavenging free radicals by synthesis of flavan-3-ols (Song et al., 2016). Os04g37820, which encodes UDP-glucuronosyl/UDP-glucosyltransferase-family protein, was reported to function as an antioxidant for detoxification (Williamson et al., 1998). In conclusion, upregulation of these 5 genes seems likely to favor scavenging of ROS and free radicals by increasing the levels of antioxidants in *idr1-1* mutant, thus aiding in *idr1-1*’s drought tolerance. There are 3 genes (Class IV), which are associated with enhancement of osmotic adjustment (Supplemental Table S4). *OsLEA18* (Os04g52110) was downregulated quite obviously in *idr1-1* mutant; LEA proteins were reported to accumulate to high levels during the late stage of seed maturation and in response to water deficit (Liu et al., 2013). However, a recent study revealed that overexpression of *ZmLEA3* in tobacco plants increased the hypersensitive cell death triggered by *pst* DC3000 and enhanced the expression of *PR1a*, *PR2* and *PR4* (Liu et al., 2013). So, it seems that downregulation of *OsLEA18* may be an indication that water contents of *idr1-1* leaves are possibly higher than those of IAPAR9 leaves, and such downregulation may be beneficial for reducing cell death. *OsASR1* (Os01g72900) and *OsASR2* (Os01g72910) each encode a ABSCISIC ACID-STRESS-RIPENING-INDUCIBLE PROTEIN, and all exhibited increased expression under control conditions. Overexpression of *OsASR1* in rice reportedly resulted in improved water regulation under salt and drought stresses, probably as a result of accumulated osmolytes and reduced transpiration rates (Park et al., 2020) All these data suggest that *idr1-1* mutant might accumulate higher levels of osmolytes to make it cope better with osmotic stress, which probably, at least in part, result from the expression changes of the aforementioned 3 genes. There are 4 genes (Class V) connected with weakening of leaf senescence, and they all showed downregulation under drought or control conditions (Supplemental Table S4). Os04g32320 encodes a glycerophosphoryl diester phosphodiesterase-family protein, and it was involved in the onset of leaf senescence (Swidzinski et al., 2002). *OsMtN3* (Os02g30910), which was downregulated in both drought and control conditions, encodes a nodulin MtN3-family protein, and it has a homologous gene in Arabidopsis, named *SAG29*; *SAG29*-overexpressing transgenic plants reportedly exhibited an accelerated senescence and were hypersensitive to salt stress (Seo et al., 2011). *OsEIL4* (Os08g39830) encodes an ETHYLENE INSENSITIVE-LIKE GENE 4 protein, and might be a positive regulator for ethylene-mediated leaf senescence (Qiu et al., 2015). *OsSAG20* (Os11g01850) is a senescence-associated gene (Mahmood et al., 2016). Collectively, these data suggest that the obviously delayed leaf senescence in *idr1-1* mutant might have come from the downregulation of such 4 genes.

There are 3 *OsGH3* genes [*OsGH3.3* (Os01g12160), *OsGH3.2* (Os01g55940) and *OsGH3.8* (Os07g40290)] (in Class VI), showing upregulation under drought or control conditions (Supplemental Table S4). GH3-family proteins catalyze IAA conjugation to amino acids, which thus leads to inactivation of IAA (Du et al., 2012). Exogenous NAA is able to induce ROS production in rice tissues, so downregulation of IAA caused by overexpression of *OsGH3.2* and the other two *OsGH3* genes (*OsGH3.3* and *OsGH3.8*) presumably contributes to reduction of ROS production (Du et al., 2012). Os01g50370, which was downregulated in *idr1-1* mutant, encodes a STE_MEKK_ste11_MAP3K.4; downregulation of this gene might help enhance resistance to water deficit through altered ABA responses (Choi et al., 2017). Overall, the above data suggest that the expression changes of these 4 genes may alter IAA levels and responses to ABA in *idr1-1* mutant, thereby resulting in attenuation of ROS production and significantly higher tolerance to drought stress. There are 6 genes with altered expression under drought or control conditions, which are connected with molecular functions shown in Class VII (Supplemental Table S4). Three genes (Os04g44060, Os01g54030 and Os02g51370) had downregulated expression under drought or control conditions, and the rest (Os09g16950, Os11g02240 and Os08g33150) exhibited upregulated expression under the same conditions. *OsPIP2;3* (Os04g44060) encodes an aquaporin. Water transport in plants has been known to greatly dependent on the expression and activity of aquaporins. However, some opposite results had been observed in OE lines of *plasma membrane intrinsic protein* (*PIP*) genes, because rapid water loss, due to increased leaf and root hydraulic conductivity, made some plants even more vulnerable to drought stress (Aharon et al., 2003). So, a general downregulation of most of the *PIP* genes under drought conditions appears to reduce water loss and to help prevent backflow of water to drying soil. *OscytME3* (Os01g54030) encodes cytosolic NADP malic enzyme 3, and restricted expression of *ZmnpNADP-ME* particularly in the guard cells of tobacco reportedly showed reduced stomatal apertures, and enhanced water use efficiency (Muller et al., 2018). *OsGRL4* (Os02g51370)*, OsCRK25* (Os09g16950), *OsCIPK15* (Os11g02240) and *Myb51-like TF* (Os08g33150) were known to be associated with stomatal closure, ROS signaling, improved tolerance to various stresses, and enhanced survival ability under stress conditions (Xiang et al., 2007; Cui et al., 2013; Zhang et al., 2013; Hu et al., 2017; Ruan et al., 2018). These data collectively suggest that alteration in expression of 6 such genes appears to contribute to increasing water retaining and survival ability of *idr1-1* mutant, thus improving its drought tolerance.

### IDR1 Physically Interacts with TUD1, and *idr1-1* Mutant Possesses Impaired BR Responsiveness

A recent study showed that rice RGA1 might genetically interact with TUD1, a U-box E3 ligase that was implicated in BR signaling; loss of function of TUD1 brought about a range of BR signaling-defective phenotypes, such as dwarfism, shortened 2nd internode, etc. (Hu et al., 2013). Our protein-protein interaction assays further proved a physical interaction between IDR1 and TUD1, suggesting a possibility that IDR1 may also take part in BR signaling (Fig. 8, A and B). In order to find out whether *idr1-1* mutants also had similar BR signaling-defective phenotypes as *tud1* mutants, we tested the responses of the *idr1-1* mutants to EBR of different concentrations by lamina joint bending and root elongation assays. The results indicated that leaf lamina joint bending in IAPAR9 seedlings was progressively increased in a BR dose-dependent manner, varying from 33° to 166° when the seedlings were treated with a range of BR concentrations (from 10^-9^ to 10^-5^ as indicated) (Fig. 8, C and D); such bending in *idr1-1* mutant seedlings also gradually increased, but the bent angles was somewhat smaller than those of IAPAR9 seedlings under the same treatment conditions, varying from 31° to 126° (Fig. 8,C and D). Besides, root elongation assay showed that elongation of primary roots of IAPAR9 seedlings were gradually inhibited with the increases in EBR concentrations (Fig. 8, E and F); by contrast, the elongation of primary roots of *idr1-1* mutant seedlings appeared less inhibited by the increasing concentrations of EBR as compared to that of IAPAR9 seedlings (Fig. 8, E and F), suggesting that such elongation of *idr1-1* mutant seedlings were of decreased responsiveness to EBR treatments with higher concentrations. These data together provide a clear demonstration that *idr1-1* mutant exhibits impaired responsiveness to higher BR concentrations than does IAPAR9 plant, which is similar to the BR-responsive phenotypes observed in *tud1* mutant (Hu et al., 2013).

**Figure 8.**
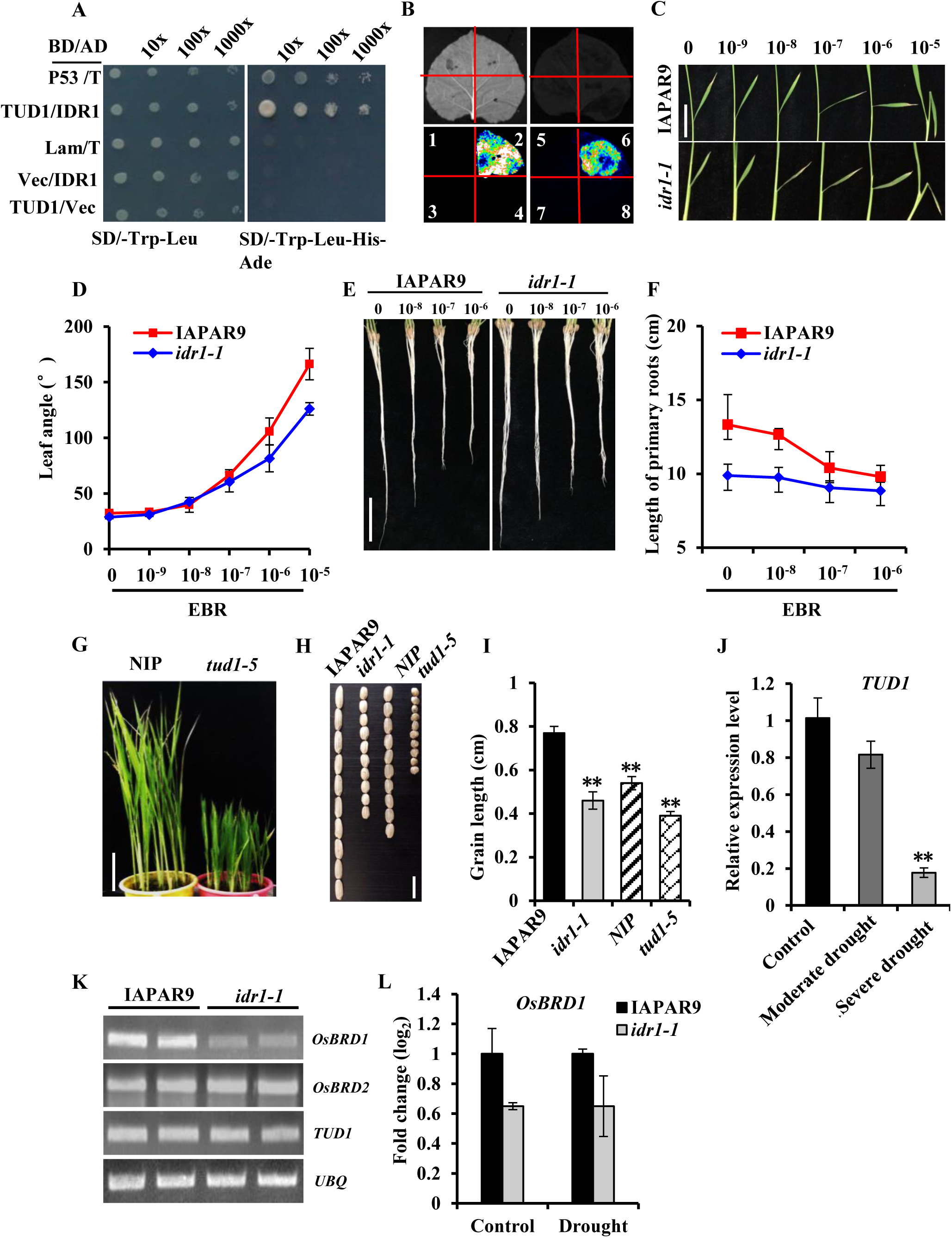
IDR1 physically interacts with TUD1, and *idr1-1* mutant exhibits impaired BR responsiveness. **(A and B)** Physical interactions between IDR1 and TUD1 in yeast cells (A) and in *N. benthamiana* leaves (B). Yeast cells (strain AH109), which was co-transfected with *BD-TUD1* plus *AD-IDR1* and other combinations of constructs as indicated, were grown on YPDA medium lacking Trp and Leu for 3 d (left panel in A) and then transferred to a stringent selection medium lacking Trp, Leu, His and adenine (SD/-Trp/-Leu/-His/-Ade) (right panel in A), and allowed them to grow for 2 days at 28°C prior to observation. In parallel, luciferase complementation image (LCI) system was used to test interactions between IDR1 and TUD1 (B). Upper panels, bright field; lower panel, dark field. 1, *AtSGT1a-nLUC* + *cLUC*; 2, *AtRAR1-nLUC* + *AtSTG1a-cLUC*; 3, *nLUC* + *AtRAR1-cLUC*; 4 and 8, *nLUC* + *cLUC*; 5, *IDR1-nLUC* + *cLUC*; 6, *IDR1-nLUC* + *TUD1-cLUC*; 7, *nLUC* + *TUD1-cLUC*. The number “2” in B indicates a positive control, and the number “4” and “8” indicate negative controls. **(C and D)** Effects of different concentrations of EBR on the degrees of inclination of the leaf lamina. Five-d-old *idr1-1* mutant and IAPAR9 seedlings grown in Hoagland solution were treated by different concentrations of EBR (0, 10^-9^, 10^-8^, 10^-7^, 10^-6^ and 10^-5^ M) for 3 days. Photographs of the second lamina joints were taken (C) and angles of the leaf lamina joint bending were measured and plotted against varying EBR concentrations (D). Data are means (±SD) for three biological replicates; for each genotype, 3 seedlings for control (0 M EBR) or each EBR concentration were used for measuring the angles of leaf lamina joint bending in each of the three replicates. Scale Bar = 5 cm. **(E and F)** Responses of primary roots of *idr1-1* mutant and IAPAR9 seedlings to different concentrations of EBR. Seeds of both the genotypes were germinated in Hoagland solution for 5 days, and the selected seedlings with uniform lengths of primary roots were treated with different concentrations of EBR (0, 10^-8^, 10^-7^ and 10^-6^ M) for 3 days. Photographs of the primary roots following treatment were taken (E), and lengths of primary roots were plotted against varying EBR concentrations (F). Data are means (±SD) for three biological replicates; for each genotype, 5 seedlings for control (0 M EBR) or each EBR concentration were used for measuring root lengths in each of the three replicates. Scale Bar = 5 cm. **(G)** Comparisons of plant heights between Nipponbare (NIP) and *tud1-5*. Seedlings of the NIP and *tud1-5* were grown in pots for 40 days prior to photography. Scale Bar = 5 cm. **(H and I)** Comparisons of grain lengths between *idr1-1* mutant and IAPAR9 plants and between *tud1-5* mutant and NIP plants. Mature seeds of IAPAR9, *idr1-1*, NIP and *tud1-5* were harvested simultaneously and photographed (H). Grain lengths were measured and compared (I). Scale Bar = 1 cm. **(J)** Relative expression levels of *TUD1* in IAPAR9 leaves that underwent moderate, severe or no (control) drought stress as revealed by qRT-PCR assays. **(K and L)** Expression levels of *OSBRD1*, *OSBRD2* and *TUD1* in *idr1-1* mutant as shown by RT-PCR assays (K). Meanwhile, *OSBRD1* transcript abundance from the transcriptome data was shown as mean log_2_ fold change ± SE (n = 3) of three independent experiments (L); values were plotted relative to the log_2_ of normalized IAPAR9 transcript abundance which was set at 1.0. RNA samples prepared from *idr1-1* mutant and IAPAR9 leaves without stress treatment (K) or leaves of both the genotypes undergoing moderate drought stress (L) were used for such assays. Data are means (±SD) for three biological replicates. Significant differences from IAPAR9 (I, L) or control (J) were determined by Student’s *t*-test: **P < 0.01, t-test.

Both *idr1-1* and *tud1-5* mutants showed growth retardation at various developmental stages compared with their corresponding wild-type controls (Fig. 8G; Supplemental Fig. S1A). Seeds produced from *idr1-1* and *tud1-5* mutants were about one-half as long as those produced from the wild-type controls (Fig. 8, H-I; Supplemental Fig. S1E). qRT-PCR assays revealed that *TUD1* expression was downregulated by moderate and severe drought stresses, which showed a similar trend in expression as observed for *IDR1* (Fig. 8J; Fig. 3A). Furthermore, RT-PCR and transcriptomic results both indicated that *BR-deficient DWARF 1* (*OsBRD1*), which is involved in BR biosynthesis, was downregulated in *idr1-1* mutant (Fig. 8K) (Hong et al., 2002). Because *TUD1* and *IDR1* were both discovered to be involved in BR signaling and both proteins physically interact with each other, it appears that *TUD1* and *IDR1* may function together in the same protein complex to help transduce BR signal through a non-canonical BR signaling pathway, and the disruption of such signal transduction may be associated with enhanced drought tolerance of *idr1-1* mutant. This notion is supported by the previous report that knockdown of BR receptor *BdBRI1* causes attenuated BR signaling, therefore rendering the plants much more tolerant of drought stress (Feng et al., 2015).

## Discussion

As a highly drought-tolerant mutant, *idr1-1* mutant has been preserved in our research group for almost 20 years. This mutant was grown every year and its drought tolerance was assessed continually in every rice growing season. It is clear that *idr1-1* mutant consistently showed very strong tolerance to drought stress under strictly controlled soil moisture regimes each year, indicating that *idr1-1* mutant is a real drought-tolerant mutant. In this study, our analyses pointed to the fact that the strong drought tolerance of *idr1-1* mutant presumably stemed from several aspects, including altered plant morphology, reduced ROS production, elevated ROS scavenging, possibly increased antioxidant contents, weakened leaf senescence, etc., all of which collectively led to the obviously increased drought tolerance as observed in *idr1-1* mutant for many years. Thus, our results provide valuable insights into the mechanisms of why loss of function of rice Gα subunit brings so strong drought tolerance to the *idr1-1* mutant.

### Morphological and Physiological Changes of *idr1-1* mutant Help Improve its Drought Tolerance by Reducing Water Loss

It had been reported that Arabidopsis GPA1 is a positive modulator of plant cell proliferation; *gpa1* mutant had reduced cell division in aerial tissues, which resulted in a reduced number of elongating cells (Ullah et al., 2001). Rice *d1* mutant defective in *RGA1* displayed marked dwarfism phenotype, demonstrating that the RGA1 serves a similar function as the GPA1 in the cell proliferation (Ashikari et al., 1999; Ueguchi-Tanaka et al., 2000). Our data also indicated that *idr1-1* mutants exhibited obvious reduction of plant height and lengths of leaves, panicles, and grains, compared to IAPAR9 plants during the whole period of growing (Fig. 1; Supplemental Fig. S1). In addition, the average leaf thickness of *idr1-1* mutants was significantly greater than that of IAPAR9 plants (Fig. 5, C and D). It is worth noting that the average root length and average root volume of *idr1-1* mutants were just slight shorter and smaller than those of IAPAR9 plants, respectively, under drought stress conditions (Fig. 5, F and G) Likewise, Arabidopsis GPA1 did not regulate cell proliferation in root meristems as well (Ullah et al., 2001), which is in agreement with our observations that the average root length of *idr1-1* mutants was slightly changed compared with that of IAPAR9 plants (Fig. 5F). Because *idr1-1* mutants had similar average root length and average root volume as did IAPAR9 plants, the significantly reduced aboveground biomass of *idr1-1* mutants must remarkably slow down water loss in such mutants compared to in IAPAR9 plants

Measurements of water potentials and leaf water loss rates revealed that *idr1-1* mutants possessed markedly higher water potentials from 10:00 to 18:00 than IAPAR9 plants, and the leaf water loss rates of *idr1-1* mutants were also decreased relative to those of IAPAR9 plants from 2 h onwards after leaf detachment (Fig. 4, A and B). *idr1-1* mutants accumulated higher levels of proline and had higher chlorophyll contents when compared to IAPAR9 plants under moderate drought conditions (Fig. 4, C and F). Moreover, *idr1-1* mutants had significantly lower MDA contents and REL (Fig. 4, D and E). Besides, *idr1-1* leaves appeared thicker than IAPAR9 leaves at the examined positions (Fig. 5, C and D). Thus, these morphological and physiological changes of *idr1-1* mutant help retain more water in tissues when the mutant plants are challenged by drought stress.

### Mutation of *IDR1* greatly Impairs Abiotic Stress- or ABA-induced Cytoplasmic and Chloroplastic ROS Burst, which is Derived from NADPH Oxidases and Chloroplasts, therefore Reducing Cell Death

It is generally known that ROS generated from chloroplasts and apoplasts are considered as major sources of ROS when plants experience abiotic stresses (drought, salinity or oxidation) or ABA treatment. There were a few reports of the involvement of GPA1 in ROS production when plants are exposed to ABA treatment or oxidative stress. Under ABA treatment, ROS production was found to be evidently impaired in *gpa1* guard cells (Zhang et al., 2011). *gpa1* mutation still allowed stomata to close to a level similar to that in Col-0 after ABA treatment, but completely abolished ABA inhibition of stomatal opening, which led to more water loss in the *gpa1* mutants (Wang et al., 2001). These data collectively suggest that GPA1 acts upstream of ROS production in the ABA signaling pathway. After MV treatment, *idr1-1* leaves showed much less ROS production than the leaves from IAPAR9 plants (Fig. 6, C and D), suggesting that chloroplast-derived ROS burst induced by MV is greatly impaired in *idr1-1* leaves. In Arabidopsis, O_3_-treated *gpa1* mutants displayed less leaf damage by excessive ROS than Col-0 control plants (Joo et al., 2005). In fact, following exposure to O_3_, *gpa1-4* mutants showed even less damage than the control plants (Joo et al., 2005). It had been observed that O_3_ treatment triggered ROS generation from chloroplasts and apoplasts of wild-type leaf cells, while the chloroplastic ROS amounts were substantially decreased and the apoplastic ROS (via NADPH oxidases) were hardly produced in the *gpa1* leaf cells. These facts together demonstrate that Gα (IDR1 and GPA1) is required for both chloroplastic and apoplastic ROS production (Joo et al., 2005).

Ethylene is able to induce H_2_O_2_ production and stomatal closure, and GPA1 is needed for the production of H_2_O_2_ induced by ethylene; *gpa1* mutants showed defects in ethylene-induced H_2_O_2_ production and stomatal closure, whereas *GPA1*-overexpressing lines showed faster stomatal closure and H_2_O_2_ production in response to ethylene (Ge et al., 2015). BR can stimulate ethylene synthesis, which ultimately leads to stomata closure through H_2_O_2_; Gα mediated BR-induced stomatal closure via promoting H_2_O_2_ and NO production (Shi et al., 2015). In *gpa1* mutant, however, ethylene or BR treatment could not effectively trigger ROS production, suggesting that GPA1 serves functions in mediating initiation and amplification for apoplastic and/or chloroplastic ROS triggered by ethylene and BR. Our results consistently showed that drought stress could also not effectively induce effective H_2_O_2_ and O_2_^-^ production in *idr1-1* mutant, suggesting that *idr1-1* mutation also impairs ROS production in rice (Fig. 6, A and B), which points to the possibility that drought stress fails to induce expression of *OsRBOH* genes or activate OsRBOHs’ activities efficiently in the *idr1-1* mutant background (Supplemental Fig. S9). In *d1* mutant, cell death in response to ethylene and H_2_O_2_ was nearly completely abolished (Steffens and Sauter, 2009). Similarly, we found that *idr1-1* mutation mitigated cell death elicited by drought stress (Fig. 6A). Therefore, ROS production and cell death triggered by ethylene, BR and drought stress are considerably impaired by Gα mutations. *d1* mutant was documented to have a similar dwarfism phenotype and impaired GA signaling as did GA-deficient mutant, which overaccumulated DELLA proteins (Ueguchi-Tanaka et al., 2000; Colebrook et al., 2014). So, the significantly increased drought tolerance in *idr1-1* mutant may partially come from DELLA overaccumulation that limits excessive ROS production (Du et al., 2012). Furthermore, our transcriptome data revealed that in *idr1-1* mutant, three *GH3* genes (in Class VI) all showed upregulation under drought or control conditions, which might facilitate decreasing IAA level, and thus contribute to reduction of ROS production (Supplemental Table S4) (Du et al., 2012).

Our transcriptome analyses revealed 941 and 120 DEGs identified under control and drought-stressed conditions, respectively (Supplemental Data Sets S1 and S2). Under drought-stressed conditions, there were 11 upregulated genes and 38 downregulated genes discovered as being regulated in expression by both *idr1-1* mutation and drought stress (Supplemental Fig. S13; Supplemental Table S3). A total of 11 genes in Class I showed altered expression in the drought and/or control conditions (Supplemental Table S4). Downregulation of 10 out of the 11 genes aids in reducing ROS production and cell death derived from hypersensitive responses (Supplemental Table S4). In addition, upregulated expression of Os01g64120 (encoding an Fd-like protein) in *idr1-1* mutant may prevent overproduction of O_2_^-^ under drought stress conditions (Hanke and Mulo, 2013). Hence, these results suggest that altered expression of the above-mentioned genes in *idr1-1* mutant favors the attenuation of ROS production and cell death under the drought-stressed conditions.

A recent study reported that when exposed to high light, *d1* mutants exhibited a greater capacity to dissipate excess irradiance relative to wild-type controls (Ferrero-Serrano et al., 2018). Therefore, D1 seems to be a regulator of photoavoidance and photoprotection mechanisms in rice (Ferrero-Serrano et al., 2018). During photosynthesis, splitting of water molecules occurs in PSII complex, and the released electrons are then transferred to the PSI complex via electron transport chain (ETC) from plastocyanin to Fd. Under stress conditions, the electrons can be transferred from Fd to O_2_ for Mehler peroxidase reaction, in which O_2_ is reduced to O_2_^-^ and H_2_O_2_ (Gururani et al., 2015); as a consequence, these reactions generate a high level of ROS, thereby causing photodamage or photoinhibition of PSII apparatus (Gururani et al., 2015). Currently, a new proposition suggests that environmental stresses do not directly cause photoinhibition but rather facilitate inhibition of PSII damage repair (Gururani et al., 2015). Moreover, stresses also aggravate ROS production within the ETC of PSII and PSI during light reactions when CO_2_ is limited and ATP synthesis is impaired (Gururani et al., 2015). ROS inhibits repair of photodamaged PSII by primarily suppressing de novo synthesis of PSII proteins, especially reaction center D1 proteins [this D1 protein is completely different from rice Gα (D1) protein] (Murata et al., 2007; Gururani et al., 2015). The damaged D1 proteins caused by photodamage are selectively degraded by a thylakoid membrane-bound metalloprotease, FtsH (Kato et al., 2009). The expression of *FtsH* was maintained by THF1 or GPA1 because *FtsH* transcript level decreased in *thf1* or *gpa1* mutant (Zhang et al., 2009). THF1 was found to physically interact with GPA1 in chloroplasts (Huang et al., 2006). When photodamaged D1 proteins were rapidly cleaved by FstH or Deg proteases into large fragments, ROS production was greatly activated, thus evoking ROS burst in the chloroplasts (Kato and Sakamoto, 2014). D1 proteins could be phosphorylated in a light-dependent manner, and such phosphorylation was able to protect D1 proteins against cleavage by FstH or Deg, which consequently diminished ROS production (Kato and Sakamoto, 2014). *gpa1* mutant showed obviously reduced chloroplastic ROS (especially H_2_O_2_) production under the conditions of oxidative stress and ABA treatment (Joo et al., 2005; Zhang et al., 2011; Ge et al., 2015), and *idr1-1* mutant also showed decreased ROS production either from apoplasts or from chloroplasts (Fig. 6), which presumably contributes to reducing damage to D1 as well as phosphorylated D1 proteins. Our qRT-PCR results demonstrated that 9 out of the 11 tested genes, such as *OsLFNR1*, *OsFd4*, *OsPC*, etc., showed higher expression levels in *idr1-1* mutants than in IAPAR9 plants under moderate and severe drought stress conditions (Supplemental Fig. S10). Besides, it is worth noting that impaired stomatal closure in *idr1-1* mutants under drought stress was helpful in reducing photorespiration (which leads to a large ROS burst) caused by CO_2_ shortage as a result of tightly closed stomata (Fig. 5, A and B). Based on the above facts, we propose a hypothesis to explain why *idr1-1* mutation results in substantial decrease in ROS production derived from PSI and PSII. For PSI, the elevated expression of the 9 genes implicated in ETC (e.g. *OsLFNR1*, *OsFd4*, and *OsPC*) in *idr1-1* mutant may contribute to successful electron transport to generate NADPH and ATP for Calvin-Benson cycle, and thereby diminish the transfer of electron to O_2_, thus leading to a marked decrease of H_2_O_2_ production from PSI in chloroplasts (Gong et al., 2020). For PSII, in the daytime, high light leads to photodamage of D1 and phosphorylated D1 proteins in wild-type plants, and such damaged D1 proteins are soon recognized and cleaved into large fragments, thus leading to excess ROS burst. However, mutation of Gα (GPA1 or IDR1) decreases FtsH level, consequently reducing cleavage on damaged D1 proteins, which therefore results in accumulation of more phosphorylated D1 or damaged D1 proteins during the daytime. At night, the phosphorylated damaged D1 proteins are converted to damaged D1 proteins through dephosphorylation, and then they are degraded progressively by FstH or Deg proteases, which does not trigger rapid ROS burst from PSII in chloroplasts (Kato and Sakamoto, 2014). Therefore, large-scale ROS burst does not appear in *gpa1* or *idr1-1* mutant when either mutant is subjected to oxidative stress and ABA treatment.

### Mutation of *IDR1* Enhances ROS-Scavenging Ability by Triggering Gene Expression of ROS-scavenging Enzymes, thus Lessening Oxidative Stress and Protecting Membrane Integrity

It is widely known that ROS scavenging is very important for drought tolerance. Plants with reduced or loss of ROS scavenging ability usually show increased drought and oxidative sensitivity (You et al., 2014). Conversely, increases of activities of ROS scavenging enzymes make a positive contribution to plant’s drought tolerance (Liu et al., 2017). Under the progressive drought treatments, the *d1* mutants reportedly exhibited obviously increased tolerance to drought stress, which is consistent with our observations that *idr1-1* mutants showed remarkably elevated drought tolerance (Ferrero-Serrano and Assmann, 2016). Our results showed that mutation of *IDR1* might have led to activated expression of 11 ROS-scavenging genes (Fig. 7). Considering that IDR1 is not only localized in the nucleus, but also at plasma membrane or cell periphery (Fig. 3H), it seems likely that IDR1 enters nucleus to repress expression of some important genes for highly efficient ROS scavenging, which is presumably one of the important causes of the significantly enhanced tolerance of *idr1-1* mutant to drought stress.

Our transcriptome data revealed that there are 10 genes (Class II), which have putative functions in ROS scavenging or detoxification, showing altered expression under drought-stressed or control conditions (Supplemental Table S4). Upregulation of the five peroxidase genes (*OsPOXs*), *OsANN3*, *OsGSTU6* and *OsLG3*, reportedly induced elevated expression of a range of ROS scavenging genes (such as *APXs*, *CATs*, *PODs*, *SODs*, etc.), which greatly contributed to eliminating excessive ROS and increasing survival rates of transgenic rice plants under drought stress conditions (Atack and Kelly, 2007; Ning et al., 2010; Xiong et al., 2018; Li et al., 2019). In addition, downregulation of *OsAAO1* in *idr1-1* mutant aided in overaccumulation of L-ascorbate for ROS scavenging (Chen et al., 2013). Hence, these data together indicate that expression changes of these genes in *idr1-1* mutant must play a very positive part in scavenging excessive ROS induced by drought stress. Moreover, our transcriptome analyses further demonstrated that there are 5 genes (Class III), which were all upregulated in drought or control conditions, associated with biosynthesis of antioxidants (Supplemental Table S4). F3H2 was reported to have a major impact on scavenging free radicals by synthesizing flavan-3-ols, and had essential roles in the survival of woody plant *Lycium chinense* when grown under extremely drought-stressed conditions (Song et al., 2016). Thus, upregulation of these 5 genes is likely to make substantial contributions to scavenging ROS and improving *idr1-1*’s drought tolerance.

Plant phospholipase D (PLD) has been shown to be involved in the action or production of several plant hormones (like ABA, ethylene, jasmonic acid, and cytokinin), cell growth, abiotic stress responses, oxidative burst etc. (Mane et al., 2007; Li et al., 2009). When plants undergo stresses, such as drought and salinity, PLDα1 hydrolyzes membrane lipids, and therefore produces a high level of PA; PA reportedly contributes to stomatal closing and reduction of transpirational water loss in the presence of ABA (Hong et al., 2008; Hong et al., 2010). *PLDα1*-suppressed or knockout plants displayed less sensitivity to ABA-induced stomatal closure and lost more water than wild-type controls (Zhang et al., 2005; Mane et al., 2007; Hong et al., 2008). Nevertheless, PLDα1 has adverse effects on plants when they experience progressively increasing drought stress, because increasing PLDα1 activities will break down membrane bilayers, and as a consequence lead to loss of cell membrane integrity, which ultimately promotes rapid water loss and leaf senescence (Hong et al., 2008). So it appears that in the early stages of drought stress or dehydration, increased expression of *PLDα1* promotes signaling of stomatal closure and decreases water loss; nevertheless, in the late stages, prolonged severe drought stress accelerates membrane deterioration, ionic leakage, lipid peroxidation and water loss (Hong et al., 2008; Li et al., 2009). Similarly, a recent study showed that Arabidopsis *PLDδ* knockout plants were much more tolerant of severe drought and displayed retarded ABA-induced leaf senescence, presumably by suppression of membrane lipid degradation (Distefano et al., 2015). It had been reported that GPA1 could bind to PLDα1, and this binding inhibited the PLDα1 activity; the interaction between PLDα1 and GPA1 could substantially decrease PLDα1 activity (Li et al., 2009). PLDα1 preferred to bind to Gα-GDP, and did not bind to Gα-GTP (Li et al., 2009). When plants were treated with ABA or drought stress, Gα-GDP was converted to Gα-GTP, and Gα was dissociated from PLDα1, which therefore relieved the inhibition of PLDα1 (Li et al., 2009). Increased expression or enzymatic activity of PLDα1 promoted signaling of stomatal closure at the earlier stages of drought stress via PA-promoted H_2_O_2_ production through RBOHs (Li et al., 2009). Gα then entered nucleus to activate and repress expression of a host of genes (like ROS-scavenging genes) as evidenced by nuclear localization of IDR1 and transcriptome data (Fig. 3H; Supplemental Data Sets S1 and S2; Supplemental Table S4). On the basis of the above facts, we put forward a hypothesis to delineate how Gα mutation abolishes PLDα1-dependent ROS production. Drought stress induces accumulation of ABA, and the ABA promotes dissociation of PLDα1 and Gα (GPA1 or IDR1). The PLDα1 then promotes generation of PA, and PA subsequently binds to RBOHs to activate their enzymatic activities (Zhang et al., 2009). Meanwhile, dissociated Gα enters nucleus to trigger expression of a group of genes, including the genes to modify ROBHs for their normal functions, and the genes to promote ROS production in chloroplasts. However, Gα seems to inhibit expression of ROS-scavenging genes, because 11 out of 13 tested genes (*OsAPX1-OsAPX2, OsAPX4-OsAPX8*, *OsCAT1*, *OsCAT3*, *OsFeSOD* and *OsPOX22.3*) were upregulated to varying extents in *idr1-1* mutant (Fig. 7B); additionally, 10 ROS-scavenging genes were also identified to be upregulated in the *idr1-1* mutant by our transcriptome analyses (Supplemental Table S4). This ultimately leads to H_2_O_2_ accumulation in cytoplasm and chloroplasts, and subsequently results in stomatal closure. By contrast, mutation of Gα fails to activate gene expression of those genes for ROBHs’ modifications as well as chloroplastic ROS production, and repress expression of ROS-scavenging genes, which eventually leads to a great decrease in ROS production in cytoplasm and chloroplasts, and a substantial increase in ROS-scavenging activity. As a result, the cytoplasmic and chloroplastic ROS levels are considerably diminished in the Gα mutant, thus abolishing or impairing stomatal closure.

### *idr1-1* Mutant Has Impaired BR Signaling, which possibly Makes Valuable Contributions to Enhanced Drought Tolerance in such a Mutant

BRs play pivotal roles in promoting cell expansion and division, regulating senescence, modulating plant responses to various environmental signals, etc. (Hu et al., 2013). When treated by BL, *gpa1* mutants showed impaired sensitivity to BL, probably suggesting that GPA1 mediates BR signaling; however, *gpa1* mutant had similar responses to ABA and ethylene as did wild-type Col-0 (Ullah et al., 2002). It had been reported that in Arabidopsis, EBR treatment initiated a marked rise in the levels of ethylene, H_2_O_2_ and NO, which were necessary for stomatal closure in wild-type Col-0 (Shi et al., 2015); nevertheless, EBR failed to close stomata of *gpa1* mutants because EBR-induced H_2_O_2_ production was greatly impaired in such mutants (Shi et al., 2015). A recent investigation showed that downregulation of *BdBRI1* resulted in reduced plant heights, shortened internodes, narrow and short leaves, reduced expression of some BR-signaling genes, and enhanced expression of some BR biosynthesis genes (Feng et al., 2015). Importantly, *BdBRI1* RNAi lines exhibited greatly enhanced drought tolerance, accompanied by highly elevated expression of some drought-responsive genes, such as *BdP5CS* (for proline synthesis), *BdCOR47*/*BdRD17*, *BdERD1* and *BdRD26* (Feng et al., 2015). In addition, triple mutations in *WRKY46*, *WRKY54*, and *WRKY70*, which were involved in both BR-regulated growth and drought responses in Arabidopsis, rendered Arabidopsis plants more tolerant of drought stress (Feng et al., 2015). Our results showed that *IDR1* was involved in rice BR responses, and mutation of *IDR1* led to impaired responses of lamina joint bending and elongation of primary roots to BR (Fig. 8, C-F); additionally, *idr1-1* mutation also brought about downregulation of *OsBRD1* (Fig. 8K). Considering the above facts, it is reasonable to conclude that the defect in BR signaling increase drought tolerance in *Brachypodium distachyon*, Arabidopsis, and rice.

*tud1* mutant showed less sensitivity to EBR treatment (Hu et al., 2013), which is similar to the situation observed in *gpa1* and *idr1-1* mutants (Shi et al., 2015). TUD1 was reported to genetically interact with D1 (Hu et al., 2013), and our protein-protein interaction assays corroborated such an interaction (Fig. 8, A and B). Both *idr1-1* and *tud1-5* mutants showed growth retardation and pronouncedly shorter grains compared with their corresponding wild-type controls (Supplemental Fig. S2; Fig. 8, G-I). These data collectively suggest a possibility that IDR1 and TUD1 both have a degree of impacts on BR signaling. However, although IDR1 and TUD1 proteins are both involved in BR signaling, they are apparently not canonical components of the BRI-mediated signaling pathway (Hu et al., 2013). Taken together, these data suggest that loss of function of IDR1 impairs BR signaling, but may have beneficial impacts on drought tolerance, probably via attenuation of ROS production and other as yet unidentified mechanisms.

### *idr1-1* Mutation Causes a Considerable Delay of Leaf Senescence, thus Enhancing Drought Tolerance

Drought-tolerant plants usually have a feature of obviously delayed drought-induced leaf senescence, and as a consequence they are able to maintain higher water contents and retain higher photosynthetic activities under drought stress conditions. Plants’ stay-green phenotype or delay in leaf senescence is known to stem from suppression of ethylene, ABA, BR, and strigolactone signal transduction or activation of cytokinin signaling, indicating that a complex signaling network regulates leaf senescence (Kusaba et al., 2013). *DET2* encodes a steroid 5α-reductase that catalyzes an early step of BR synthesis, and *det2* mutant showed a stay-green phenotype (Chory et al., 1994). *bzr2*/*bes1*, a transcription factor positively regulating BR signaling, also showed a stay-green phenotype (Yin et al., 2002). These and other lines of evidence strongly suggest that the disruption of either BR biosynthesis or signaling remarkably delays leaf senescence. *gpa1* mutant showed attenuated leaf senescence, delayed chlorophyll degradation under stress conditions (Urano et al., 2014). *PLDα1*-deficient transgenic plants were reported to show slower rates of leaf senescence (Fan et al., 1997). Our results indicated that in upland fields, leaves of *idr1-1* plants remained green and became hardly senescent even after full seed maturity, indicating that *idr1-1* plants have much delayed leaf senescence and breakdown of leaf proteins as well as membranous structures even following seed maturity. Upon 3-d-darkness induction, *idr1-1* leaves displayed delayed leaf senescence and weakened ROS and O_2_^-^ production (Supplemental Fig. S8). As *OsBRD1* transcript level was downregulated (Fig. 8K) and BR signaling was impaired in *idr1-1* mutants (Fig. 8, C-F), it is possible that the severely delayed leaf senescence occurring in *idr1-1* mutants is a result of impaired BR biosynthesis or signaling. Alternatively, impaired functions of PLDα1 may make positive contributions to such delay in leaf senescence.

Transcriptome analyses revealed that there are 4 genes (Os04g32320, *OsMtN3*, *OsEIL4* and *OsSAG20*) (Class V) associated with weakening of leaf senescence; all of them were downregulated under drought or control conditions (Supplemental Table S4). The proteins encoded by the aforementioned genes take part in initiation of leaf senescence, ethylene-mediated leaf senescence, or hormone-promoted leaf senescence. Therefore, the decreases in expression of such genes might have contributed to delay in leaf senescence under drought stress conditions.

### *idr1-1* Mutation Increases Gene Expression of Drought-tolerant Genes, therefore Improving Survival Ability of the Mutant under Drought Stress Conditions

Data from other studies revealed that RGA1 regulated expression of an abundant group of genes. For example, in *rga1* mutant, there were 626 genes, which separately responded to heat (522), drought (101), salt (37), and cold (15) stresses, found to be changed in gene expression levels (Jangam et al., 2016). Among these genes, a few genes were implicated in biosynthetic pathways of osmoprotectants, such as polyamine, glycine-betaine, proline, and trehalose (Jangam et al., 2016). These data indicate that RGA1 does indeed participate in regulation of expression of the genes associated with abiotic stresses. Our transcriptome data demonstrated that there are 4 genes (Class IV), which are associated with enhancement of osmotic adjustment, displaying altered expression levels in drought and/or control conditions (Supplemental Table S4). Downregulatio of *OsLEA18* seems to help weaken cell death; upregulation of *OsASR1* or *OsASR2* are linked to accumulation of osmolytes (Supplemental Table S4) (Desikan et al., 2006; Du et al., 2020; Park et al., 2020). Hence, the alteration of expression of these genes in *idr1-1* mutant might confer better osmotic adjustment to such a mutant, and as a consequence improve the survival ability of the *idr1-1* mutant under drought stress conditions. There are 7 genes (Class VII) showing expression changes under drought or control conditions, which participate in increases of water retaining, survival ability, and drought tolerance (Supplemental Table S4). Decreased expression of *OsPIP2;3* under drought conditions was thought to aid in reducing water loss by preventing backflow of water to drying soil. Downregulation of *OscytME3* or *OsGRL4* appears to result in reduced stomatal apertures, and enhanced water use efficiency (Hu et al., 2017; Muller et al., 2018). In addition, upregulation of *OsCRK25*, *OsCIPK15* or *Myb51-like TF* contributes to improving the damaging effects of cold, drought, and salt stresses, and increasing water retaining and survival ability of *idr1-1* mutant under drought-stressed conditions.

Altogether, in this study, we detailed the effects of the *idr1-1* mutation on the drought tolerance, and found that loss of function of IDR1 very obviously enhanced tolerance to drought stress in rice. Mutation of *IDR1* led to expression changes of a few dozen genes that were related to ROS production, ROS scavenging, antioxidation, osmotic adjustment, leaf senescence, etc., suggesting that improved drought tolerance in *idr1-1* mutant cannot be simply attributable to its dwarfism phenotype, but rather results from multiple layers of regulations occurring at morphological, physiological and molecular levels. Gα mutation in Arabidopsis and rice affects multiple signaling pathways, suggesting that GPA1 serves to integrate quite a few internal and external signals, which makes it a good material to study the molecular mechanisms of a little important signaling, such as GA, BR, ABA, ET, phospholipids, etc. *idr1-1* mutants showed significantly lower yield loss under severe drought stress conditions, implying that it can be used directly in breeding to improve drought tolerance of rice cultivars. In fact, our breeding practices have demonstrated that introduction of *idr1-1* mutation to a few cultivars does indeed improve drought tolerance of such cultivars, but does not cause significant decreases of crop yields because some progeny carrying *idr1-1* mutation have no noticeable reductions in panicle and seed lengths compared to the corresponding parental cultivars. Therefore, the *idr1-1* allele has great potential for applications in rice breeding to generate elite rice cultivars with significantly increased drought tolerance in the near future.

## Materials and methods

### Plant materials

The upland rice cultivar IAPAR9 was used for this experiment as a starting material. The *idr1-1* mutant was derived from a M2 population of IAPAR9 M1 seeds undergoing ^60^Co γ-ray (250 Gy) radiation. Other upland rice cultivars, such as HD297, HD385, IRAT109, HD119, etc., were used as male parents for crossing with the *idr1-1* mutants to generate F_2_ populations for mapping *IDR1* locus.The *tud1-5* was a newly discovered mutant allele in the Nipponbare background as described previously (Hu et al., 2013). All important primers used in this study had been listed in Supplemental Table S5.

### Drought stress treatments

For drought-stressed treatments in pots (20 cm in diameter and 15 cm in depth), seedlings at a V6 (i.e. when the sixth leaf had a visible “collar” at the base of the leaf) developmental stage were subjected to drought stress by withholding water. When most of the IAPAR9 leaves were dried but the *idr1-1* mutant was affected moderately, we defined such a scale as the severe drought stress conditions; at that time soil moisture was close to 5% [determined by a precise moisture measurement (PMM) with a TRIME-PICO64 TDR Technology instrument (IMKO Micromodultechnik GmbH Company, Ettlingen, DE, Germany)]. After severe drought stress, the plants were rewatered, and survival rates of *idr1-1* mutant and IAPAR9 plants were analyzed.

For drought-stressed treatments in concrete tanks (length × width × depth: 3.2 × 2 × 2 m), all tanks were irrigated every three days until the onset of drought treatment after sowing seeds. Soil moisture content after full irrigation was approximately 50% (v/v) at a depth of 20 cm as determined by the PMM. The degrees of drought stresses in this experiment were defined as mild, moderate, and severe drought stresses when the soil moisture contents decreased to 15%, 10%, and 5%, respectively, as measured by the PMM. For example, in order to create severe drought stress, water withholding was not stopped until leaves of the *idr1-1* mutant and IAPAR9 plants displayed marked differences in leaf damage, and the bottom line of soil moisture was about 5% as determined by the PMM. Then the tanks were fully irrigated and the plants were allowed to recuperate. Survival rates were calculated one week after irrigation. Non-drought control tanks were kept irrigated every 3 days. Thirty plants were measured in each replicate and all the experiments had three biological replicates. All of the measurements were completed in one day.

### Map-based cloning of *IDR1*

To generate mapping populations, *idr1-1* mutant females were separately crossed with 10 upland rice cultivar males, including IAPAR9, HD297, HD385, IRAT109, HD119, etc. The resulting F_2_ progeny derived from such crosses were used to test segregation ratios of drought-tolerant to drought-sensitive individuals. To select drought-tolerant individuals, water was withheld from 4-week-old F_2_ seedlings for drought treatment. Finally, a total of 160 F_2_ individuals with the tolerance to moderate drought stress were selected for primary mapping using simple sequence repeat (SSR) markers (http://www.gramene.org/microsat/ssr.html) distributed across all the 12 chromosomes. Fine mapping was performed using 3800 segregants selected from 20310 F_2_ plants. The new Indel markers were developed using RiceVarMap database (http://ricevarmap.ncpgr.cn/). CAPs and dCAPs markers were designed by comparing sequences between IAPAR9 and HD119 (Yang et al., 2016). To find genomic lesions of the candidate genes, genomic DNA of the four candidate genes located between markers IM63 and IM59 from *idr1-1* mutant and IAPAR9 plant was PCR-amplified, and then the PCR products were sequenced and analyzed by DNAMAN software.

### Generation of transgenic lines

To generate *IDR1* complementation and OE lines, a 4020-bp genomic DNA sequence plus 2500-bp 5’-terminal region upstream of ATG codon and one 1173-bp ORF sequence of *IDR1* were cloned into the binary vector pCAMBIA1305 and pCAMBIA1300, respectively; the *IDR1* ORF sequence was driven by a maize ubiquitin promoter. Both the resulting constructs were separately transformed into calli derived from mature *idr1-1* seeds through the Agrobacterium (EHA105)-mediated transformation. Rice transformation was performed according to the method described previously (Hiei et al., 1994).

### Subcellular localization of IDR1

To investigate the subcellular localization of IDR1, the 1170-bp ORF sequence of *IDR1* without the stop codon was cloned into the *pCaMV35S:GFP* binary vector to generate *pCaMV35S:IDR1-GFP* construct. This construct was subsequently co-transformed with a construct carrying marker gene *plasma membrane intrinsic protein 2A* (*PIP2A-RFP*) (for plasma membrane localization) or *MADS-box transcription factor 3-like protein* (*MADS3L-RFP*) (for nucleus localization) into *Nicotiana benthamiana* leaf cells by using Agrobacterium-mediated transformation methods. The transformed cells were then examined under a confocal fluorescence microscope (Zeiss LSM 710, Germany) after 48-72 h of incubation. The excitation/emission wavelengths were 488 nm/500–550 nm for GFP, and 543 nm/620– 630 nm for mCherry.

### Measurements of morphological traits

For observation of stomata, 5-week-old leaves were harvested after severe drought stress treatment in concrete tanks, and stored in 2.5% glutaraldehyde stationary liquid (pH 7.3). The leaves were then transferred to a solution containing 1% osmic acid for 1 h before photography. Scanning electron microscopy (SEM) was used to examine stomata as previously described (Luo et al., 2013). Stomata were imaged by an Olympus BX53 microscope with IpExp60C software. Stomatal aperture was expressed as average width of a number of stomata and the apertures were measured by using the free software ImageJ (National Institutes of Health) (Ling et al., 2017). To measure thickness of IAPAR9 and *idr1-1* leaves, flag leaves of both genotypes were cut into about 1 × 1 mm by hand. The thickness of transverse sections of IAPAR9 and *idr1-1* leaves at the position in the proximity of upper portions of the main veins were observed and measured with a light microscope with CCD camera. For observation of deposition of wax crystals, the flag leaves of 30-d-old seedlings were cut into pieces of 1 × 1 cm. After being quickly frozen with liquid nitrogen, cuticular wax crystals of *idr1-1* mutant and IAPAR9 leaves were viewed by scanning electron microscope (S-3000N, Hitachi). To investigate root traits, we planted one seedling in each PVC tube, which contained mixed soil [soil/vermiculite/peat, 1:1:1 (v/v/v)]. The soil moisture was about 50% before drought treatment. When the seedlings reached a V8 developmental stage, water was withheld for drought stress. When the soil moisture decreased to about 10% (moderate drought stress), where the IAPAR9 plants began to flower, whole roots of both the genotypes were washed and the clean roots were used for measuring root traits

### Measurements of physiological traits

After moderate drought treatment in the concrete tanks, leaf water potentials of *idr1-1* mutant and IAPAR9 leaves were determined with a Pressure Chamber Instrument (Model 600) (PMS Instrument Company, Albany, OR, USA), and manipulations of the instrument were performed as described by the manufacture. Thirty leaves from each genotype were selected randomly, and water potentials of the leaves were measured at 6, 10, 14, and 18 o’clock, respectively. For measurement of water loss rate, detached leaves from *idr1-1* mutant and IAPAR9 plants as well as the three complementation lines were weighed immediately and recorded as the initial weights, and then they were incubated at room temperature to lose water gradually. Water loss was monitored by weighing these leaves at the indicated time points. Proline contents were measured as previously reported (Verslues, 2010). Briefly, leaf tissues were cut into pieces and extracted with 3% sulfosalicylic acid, and then the extract was analyzed using a spectrophotometer (UV762/762PC) at 520 nm. For each genotype, 30 plants were used for measuring the leaf proline contents in each of the three replicates. MDA was extracted as previously described (Meir et al., 1992). Briefly, 0.5 grams of leaf tissues were ground into homogenate in 5 mL solution that contained 20% trichloroacetic acid (TCA) and 1.5 mM EDTA. MDA was determined using the thiobarbituric acid (TBA) test as described in the reference. Chlorophyll contents were measured by the SPAD method (Wu et al., 2015). Briefly, total chlorophyll in the leaves was extracted with 80% acetone. The extract was analyzed using a spectrophotometer (UV762/762PC) at 470 nm. Besides, leaf electrolyte leakage was measured as described previously (Zhu and Xiong, 2013). For measurement of POD activities, briefly, 0.2 grams of leaves were ground to homogenate by adding 2 mL of PBS solution, and then the homogenate was centrifuged at 12,000 g for 20 min at 4°C. The supernatant was used for extracting POD enzyme. Peroxidase activity of POD was determined according to the method described by Kwak et al. with minor modifications (Kwak et al., 1995). The reaction mixture (in a total volume of about 200 mL) contained 0.2 M PBS (200 mL, pH 6.0), 30% (v/v) H_2_O_2_ (0.112 mL), and 0.076 mL guaiacol as substrates. Thirty microliter of enzyme extract was mixed with 3 mL of reaction mixture for each reaction. The oxidation of guaiacol was monitored and the absorbance was measured at 470 nm every 30 s.

### Measurements of responsiveness of *idr1-1* mutants to various experimental treatments

For glucose treatment, *idr1-1* mutant and IAPAR9 seeds were placed in petri dishes with different concentrations of glucose and cultured in greenhouse for 5 days. Seeds were considered to be germinated when the radicles completely emerged from husks and then photographs were taken (Chen et al., 2006). For salt treatment, 2-week-old *idr1-1* mutant and the IAPAR9 seedlings were treated with 150 mM NaCl until the salt-sensitive phenotype appeared. For MV treatment, roots of 2-week-old *idr1-1* mutant and IAPAR9 seedlings were submerged into 30 μM MV solution for 3 days. Detached leaves from both the genotypes were stained with DAB and photographed. For examination of responsiveness to EBR, 5-d-old *idr1-1* mutant and IAPAR9 seedlings grown in Hoagland solution were treated with different concentrations of EBR for 3 days in the dark. Photographs of the second lamina joint were taken and the angles of leaf lamina joint bending were measured and plotted against different EBR concentrations. For root elongation inhibition assay, 5-d-old *idr1-1* mutant and IAPAR9 seedlings were treated with EBR of different concentrations for 3 days. Lengths of the primary roots were measured and plotted against different EBR concentrations (Wang et al., 2006). For test of darkness-induced leaf senescence, detached leaves of *idr1-1* mutant and IAPAR9 plants were incubated in the dark for 3 days. DAB and NBT staining of the leaves were performed and such leaves were then photographed.

### RNA extraction and gene expression analyses

Generally, 150 mg of tissues from plant materials in each experiment were used for total RNA isolation by TRIpure Reagent (Aidlab, Beijing, China). Then the first-strand cDNA synthesis was primed with an oligo (dT) primer by using a PrimeScript™ RT reagent Kit with gDNA Eraser (Perfect Real Time) (Takara, Dalian, China) according to the manufacturer’s instructions. The resulting cDNA was then used for quantitative real-time PCR (qRT-PCR) by using a Hieff qPCR SYBR Green Master Mix (Low Rox Plus) (YEASEN, Shanghai, China) in an Applied Biosystems Q5 Real-time PCR System with primers corresponding to each detected gene. The results were normalized using *Ubiquitin5* as an internal control. RT-PCR (qRT-PCR) amplification was carried out as follows: 95°C for 5 min, followed by 18-40 amplification cycles [95°C for 30 (5) s, 55 (60)°C for 30 s, and 72°C for 30 (0) s]. The primers were listed in Supplemental Table S5. To analyze expression levels of *IDR1* under different abiotic stresses, 2-week-old IAPAR9 seedlings grown in 1/2 MS solution were subjected to different treatments with 20% (w/v) PEG 6000, NaCl (150 mM), cold (4°C), heat (40°C) or ABA (50 μM), respectively. Leaves were harvested at 0, 1, 4, 8 hours after treatment (Xiong et al., 2018). To examine expression levels of *IDR1* in different tissues and of *OsBRD1* and *OsBRD2* as well as *TUD1*, total RNA was isolated from 4-week-old *idr1-1* mutant and IAPAR9 seedlings, and then subjected to qRT-PCR assays. To check expression levels of ROS-scavenging and *OsRBOH* genes, total RNA was isolated from leaves of both *idr1-1* mutant and IAPAR9 plants undergoing moderate, severe or no stress, and used for qRT-PCR assays (Ning et al., 2010; Lu et al., 2020).

### Detection of ROS accumulation and cell death

For measurement of H_2_O_2_ accumulation in leaves, rice leaves were treated with DAB staining solution (1 mg/mL, pH 3.8) for 12 h in the dark. After 2-3 rinses, the leaves were decolorized with 100%, 90%, 80%, and 70% (v/v) alcohol each for 10 min at 100°C to gradually eliminate chlorophyll thoroughly. At least 5 leaves from different plants were analyzed for each experiment, and the representative images were shown (Xiong et al., 2018). O_2_^-^ was detected by NBT staining. Briefly, rice leaves were treated with NBT staining solution (1 mg/mL) for 12 hours in the dark and then they were decolorized. Trypan blue staining was performed following a protocol reported previously (Xiong et al., 2018). Leaves were treated with 0.1% Trypan blue solution for 12 h at room temperature in the dark followed by decolorization. CeCl_3_ was used for localizing and visualizing H_2_O_2_ at the subcellular level by transmission electron microscopy (TEM) as described previously (Zhang et al., 2010). Briefly, leaf tissues undergoing drought stress were excised to pieces (1-2 mm^2^) and incubated in 5 mM CeCl_3_ for 1 h at 4°C. Then the leaf sections were subjected to a series of manipulations, i.e. fixation, wash, postfixation, dehydration and embedment. The resulting blocks were then sectioned (70-90 nm) on a Reichert-Ultracut E microtome (Reichert-Jung, Leica, Bensheim, Germany) and mounted on uncoated copper grids (300 mesh). Sections were examined by using a TEM at an accelerating voltage of 75 kV (Zhang et al., 2010).

### mRNA-Seq analyses

Leaves of *idr1-1* mutant and IAPAR9 plants were harvested and used for mRNA-seq after moderate drought treatment. To prepare samples for mRNA-seq, 5-week-old *idr1-1* mutant and IAPAR9 plants were grown in concrete tanks for nonstressed or drought-stressed treatment. Samples were collected when soil moisture content was decreased to 10%. Total RNA was extracted from seedling using RNAiso Plus (Takara, D9108B) for mRNA-seq. mRNA-seq was carried out according to procedures provided by the sequencing company (BGI, Beijing, China). The trimmed high-quality clean data were aligned to the entire Col-0 reference genome (TAIR10) by using Hisat2 software (Kim et al., 2019), whereby parameters were set as default parameters. Subsequently, the reads were counted by using softwares FeatureCounts and Subread (Liao et al., 2013). DESeq2 was then used to analyze the gene expression levels of differentially expressed genes; meanwhile, the critical thresholds for screening for differentially expressed genes were P-value < 0.05 and the log_2_ fold change > 2 (Love et al., 2014).

### Yeast two-hybrid and LCI assays

Protein-protein interaction assays were performed as described previously (Wang et al., 2018). Briefly, for yeast two-hybrid assay, the entire coding sequences of *IDR1* and *TUD1* were in-frame cloned into pGADT7 (AD) or pGBKT7 (BD) vectors to generate fusion constructs, respectively. Then the *pGADT7:IDR1* and *pGBKT7:TUD1* constructs were co-transformed into yeast strain AH109. Later on, the co-transformed yeast cells were first spotted onto a synthetic dropout medium without Leu and Trp (SD/-Leu/-Trp) to grow for 3 days and then transferred to a stringent selection medium [synthetic dropout medium lacking Trp, Leu, His and adenine (SD/-Trp/-Leu/-His/-Ade) to grow for an additional 2 days at 28°C prior to observation. For LCI assays, the coding sequences of *IDR1* and *TUD1* were cloned into *pCAMBIA1300-nLUC/cLUC* vectors. LCI assays were performed by transiently co-expressing both constructs in tobacco leaves through Agrobacterium-mediated infiltration. Three days after agroinfiltration, luciferase activity was detected with a CCD camera (Tanon, Shanghai, China).

## Accession number

All the mRNA-seq data used in this article have been deposited in GenBank as accession number SRP273943.

## Acknowledgments

Data analyses were supported by the high-performance computing platform of Bioinformatics Center, Nanjing Agricultural University.

## Supplemental materials

**Supplemental Figure S1.** i*d*r1*-1* mutants display noticeably reduced elongation in major agronomic traits.

**Supplemental Figure S2.** i*d*r1*-1* mutants show enhanced sensitivity to glucose and NaCl treatments.

**Supplemental Figure S3.** Tests of drought tolerance of *idr1-1* mutant and other relevant lines.

**Supplemental Figure S4.** Tests of drought tolerance of *IDR1* complementation lines.

**Supplemental Figure S5.** Performance of panicles and seed setting rates of *idr1-1* mutant under severe drought stress conditions.

**Supplemental Figure S6.** Accumulation of cuticular wax crystals on flag leaves.

**Supplemental Figure S7.** Performance of root traits under drought-stressed conditions.

**Supplemental Figure S8.** Darkness-induced senescent performance in *idr1-1* mutant leaves.

**Supplemental Figure S9.** Transcript levels of a few *OsRBOH* genes in *idr1-1* mutant.

**Supplemental Figure S10.** Expression changes of a dozen genes involved in PET chain.

**Supplemental Figure S11.** Induction of expression of two marker genes, *OsRAB16C* and *OsRAB16A*.

**Supplemental Figure S12.** Materials of *idr1-1* mutant and IAPAR9 plants for transcriptome profiling.

**Supplemental Figure S13.** Analyses of the genes regulated by IDR1.

**Supplemental Table S1.** Phenotype segregation analyses of the F_2_ progeny.

**Supplemental Table S2.** Information of the genes detected by (q)RT-PCR assays.

**Supplemental Table S3.** A list of genes affected by both *idr1-1* mutation and drought stress.

**Supplemental Table S4.** Seven classes of genes exhibited significantly differential expression.

**Supplemental Table S5.** Primer sequences used in this study.

**Supplemental Data Set S1:** Differentially expressed genes between nonstressed *idr1-1* mutant and IAPAR9 plants.

**Supplemental Data Set S2:** Differentially expressed genes between drought-stressed *idr1-1* mutant and IAPAR9 plants.

## Notes

**Funding information** This work was supported by grants from the National Key Research and Development Program of China (2016YFD0100604) (to H. L.) and Foundation for Jiangsu Postdoctoral Fellowship (2019K146) (to Q. W.).

